# Early demyelination by off-target complement injury in a mouse model of neuromyelitis optica

**DOI:** 10.64898/2026.03.13.711636

**Authors:** Selin Kenet, Marina Herwerth, Shahrzad Askari, Katharina Eichenseer, Klaus Dornmair, Xuebin Qin, Christine Stadelmann, Jeffrey L. Bennett, Bernhard Hemmer, Thomas Misgeld

## Abstract

Neuromyelitis optica spectrum disorder (NMOSD) is an autoimmune CNS disease characterized by serum antibodies targeting astrocytes for complement-mediated lysis. NMOSD lesions show not only astrocyte loss but also early demyelination and oligodendrocyte injury – histological hallmarks that converge with those of other primary demyelinating conditions. How pathology spreads to cause demyelination, particularly in early lesions, remains unclear. Using spinal cord imaging in an acute mouse NMOSD model, we directly observe the spread of pathology from astrocytes to oligodendrocytes in evolving lesions. This spread is characterized by initial calcium dyshomeostasis in oligodendrocytes followed by delayed, non-lytic cell death. Oligodendrocyte death is driven not by astrocyte loss per se but by spill-over of soluble complement proteins, as oligodendrocytes can be cell-autonomously preserved by the cell-type-specific expression of the complement inhibitor CD59. Our findings explain the convergence of glial pathology in antibody-mediated CNS autoimmunity and point towards new approaches to prevent secondary glial injury.

## Introduction

Demyelination is a common hallmark of autoimmune conditions affecting the central nervous system (CNS), including neuromyelitis optica spectrum disorder (NMOSD). NMOSD primarily affects the optic nerve and spinal cord and is often associated with poor clinical recovery (Wingerchuk *et al*., 2015; Wingerchuk & Lucchinetti, 2022). The majority of NMOSD patients harbor serum antibodies against an astrocytic antigen, aquaporin 4 (AQP4-IgGs), which is expressed in the CNS neuropil on pial and perivascular astrocyte endfeet (Lennon *et al*., 2004, 2005). Rodent NMOSD models that involve local or systemic antibody delivery have established the pathogenic potential of AQP4-IgGs (Bennett *et al*., 2009; Bradl & Lassmann, 2014; Duan & Verkman, 2020). These autoantibodies are assumed to directly target a complement-mediated attack on astrocytes (Hinson *et al*., 2007; Misu *et al*., 2007; Ratelade *et al*., 2013; Roemer *et al*., 2007). Despite the apparent specificity of the initial attack, injury to other cell types is also prominent in NMOSD lesions (Lucchinetti *et al*., 2002; Misu *et al*., 2013; Parratt & Prineas, 2010; Roemer *et al*., 2007) and replicated in rodent models, including axonal damage and demyelination (Duan *et al*., 2018; Saadoun *et al*., 2010; Tradtrantip *et al*., 2017; Wrzos *et al*., 2014). Thus far, most studies of how human and experimental NMOSD pathology evolves have focused on advanced lesions, which are already aggravated by infiltration of peripheral immune cells (Wrzos *et al*., 2014). This has limited insight into the nature of immediate cellular events following the primary complement-mediated astrocytic injury, such as the mechanisms triggering demyelination and oligodendrocyte loss, processes that experimental systems suggest unfold within 24 hours (Wrzos *et al*., 2014).

Understanding the initial events in lesion formation is particularly relevant in NMOSD, as, unlike other autoimmune conditions such as multiple sclerosis (MS), where damage accrues incrementally, even a single NMOSD attack can result in permanent disability (Berthele *et al*., 2023). Preventing pathology spread based on a pathomechanistic understanding in this acute phase could thus be valuable as a first-line intervention, even before immune-modulating therapies targeting complement or antibody availability (Kumpfel *et al*., 2024) can be instituted. Notably, secondary pathology is not unique to NMOSD. Similarly, lesions in MS or myelin oligodendrocyte glycoprotein antibody disease (MOGAD), which are likely driven by oligodendrocytic autoimmune targets (Liu *et al*., 2017; Owens *et al*., 2023; Takai *et al*., 2026; Takai *et al*., 2020), exhibit – to varying degrees – convergence on pan-glial injury and axonal damage. Therefore, insights from NMOSD and its models into the mechanisms underlying interglial pathology spread might yield general lessons for a broader range of CNS autoimmune conditions.

Previous studies have suggested a range of potential mechanisms for secondary demyelination in NMOSD. One possibility is oligodendrocyte dysfunction secondary to astrocyte loss. Astrocytes provide essential metabolic and homeostatic support for other cells of the CNS, including the maintenance of potassium and water homeostasis, trophic support, or glutamate buffering (Marignier *et al*., 2010; Masaki, 2015; Richard *et al*., 2020; Sharma *et al*., 2010; Wrzos *et al*., 2014). Other mechanisms may contribute during chronic disease phases. These include nonspecific immune injury of oligodendrocytes caused by the infiltration of granulocytes and their cytotoxic secretions (Lucchinetti *et al*., 2002; Misu *et al*., 2013), by cytokines (Fujihara *et al*., 2020), or complement components that cause collateral damage (Duan *et al*., 2018; Tradtrantip *et al*., 2017).

Here, we elucidate the immediate cellular events in experimental NMOSD white matter lesions that follow a primary autoantibody/complement-mediated attack on astrocytes using in vivo imaging in mice. We reveal structural and functional changes in astrocytes and other glial cells, including oligodendrocytes, using a spinal model of NMOSD that we previously developed to characterize mechanisms of axonal injury (Herwerth *et al*., 2016; Herwerth *et al*., 2022). Our in vivo observations, buttressed by cell-type-specific genetic interventions, reveal the key contribution of activated complement proteins, specifically membrane attack complex (MAC) proteins derived from targeted astrocytes, in causing myelin and oligodendrocyte injury. In contrast, AQP4-IgG-mediated astrocyte loss per se is not sufficient for oligodendrocyte damage. This MAC-dependent astrocyte-oligodendrocyte injury seems to be specific (as other glial cells, such as microglia and oligodendrocyte precursor cells, as well as axons, are not affected by this process), mechanistically distinct (as oligodendrocyte death appears not to be lytic), and commutative (in that targeting oligodendrocytes in an MOGAD-related approach can also damage astrocytes in our model). Importantly, this pattern of off-target oligodendrocyte injury offers the prospect of rescuing sublethally injured oligodendrocytes, which could prevent secondary immune activation and provide an additional cellular source of remyelination (Duncan *et al*., 2018).

## Results

### Oligodendrocytes are injured within hours of astrocyte loss in experimental NMOSD lesions

Secondary myelin damage and oligodendrocyte injury have been observed in both human and experimental lesions of AQP4-IgG^+^ NMOSD (Carnero Contentti & Correale, 2021). To explore the early signs of secondary glial damage and the potential mechanisms underlying the intercellular spread of glial injury in experimental NMOSD, we performed in vivo multiphoton imaging in an acute spinal model that we previously established (Herwerth *et al*., 2016; Herwerth *et al*., 2022). We simultaneously imaged transgenically labeled astrocytes and oligodendrocytes (*ALDH1L1*:GFP; *PLP*:CreERT x *CAG*:fl tdTom mice; (Doerflinger *et al*., 2003; Madisen *et al*., 2010; Yang *et al*., 2011) following the local application of AQP4-IgG^+^ patient plasma (AQP4-IgG) combined with a human complement source to the dorsal spinal surface for up to 8 hours (**Supp. Video 1**). This led to the lytic depletion of astrocytes, starting within the first hours of treatment (astrocyte loss t_50%_ = 78.0 ± 8.3 min; **Fig. 1A-C**). Oligodendrocyte injury developed with a delay (oligodendrocyte injury t_50%_ = 340.6 ± 29.6 min; **Fig. 1A-D**), starting after most of the astrocytes were already depleted (**Fig. 1C**). Overt loss of oligodendrocytes was preceded by the development of a grainy pattern of fluorescent particles in neuropil, which on closer inspection proved to be beaded oligodendrocyte processes and internodes (**Supp. Video 2**). Progressively, the oligodendrocyte somata also exhibited a swollen, spheroidal shape, with increasingly prominent nuclear sparing (Olig_PN_; **Fig. 1B**, pink arrowheads at 280, 360, and 430 min). Finally, the swollen oligodendrocytes lost fluorescence (Olig_Loss_; **Fig. 1B**), with an average delay of 147 minutes (**Fig. 1D**). No glial pathology was observed under control conditions following the application of healthy donor plasma combined with the human complement source (Ctr-IgG, **Fig. 1A**).

**Figure 1.**
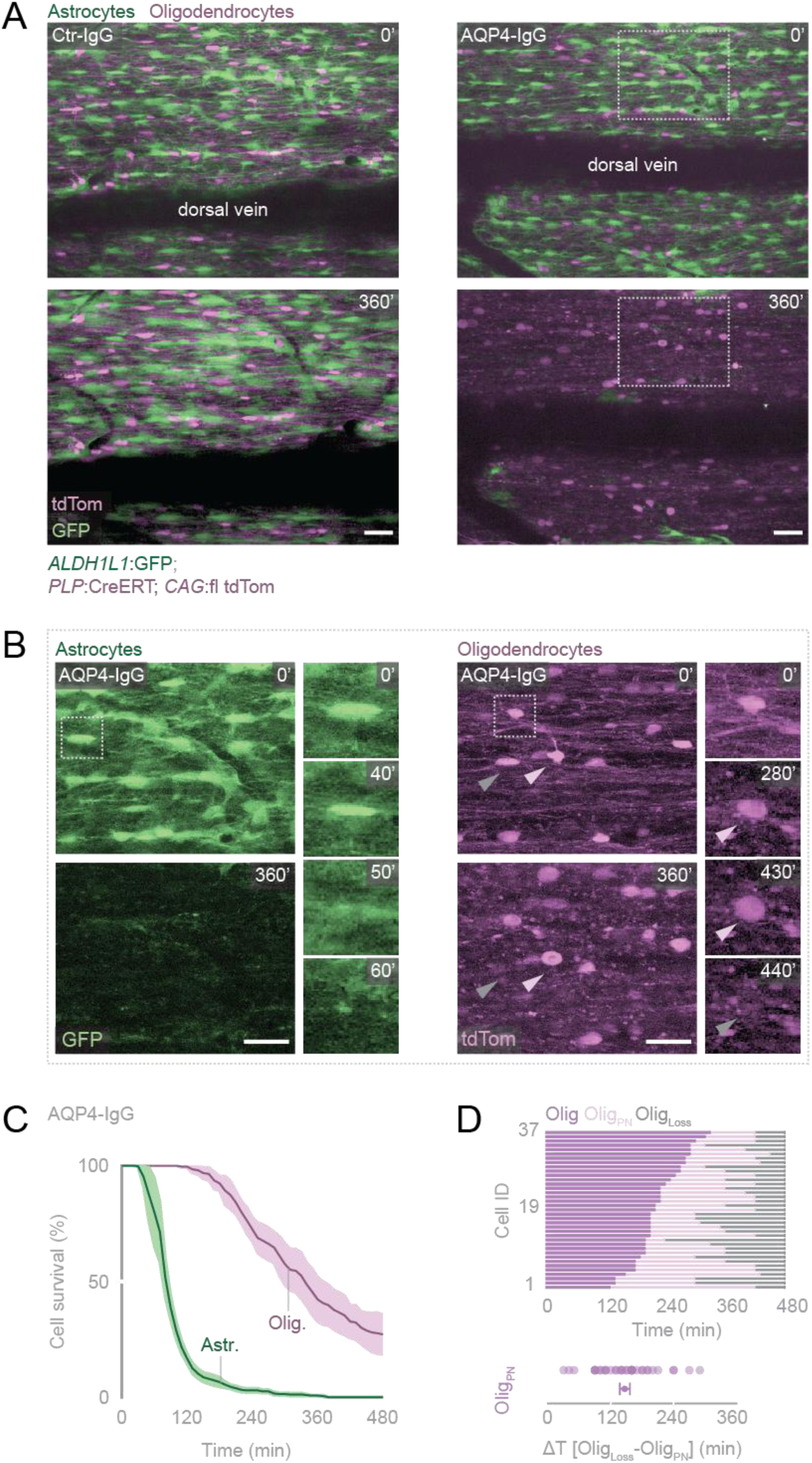
Oligodendrocyte injury follows astrocyte loss in experimental NMOSD lesions. **A.** Spinal cord in vivo imaging (maximum intensity projections) of experimental NMOSD lesions induced by AQP4-IgG and complement in transgenic mice (astrocytes, green; oligodendrocytes, magenta). Swift and global astrocyte loss followed by delayed oligodendrocyte damage, initially characterized by swollen somata with darker (“prominent”) nuclei (**right**). Astrocytes and oligodendrocytes remain unaffected in control experiments (Ctrl-IgG and complement application; **left**). Areas indicated with the white dashed boxes are magnified in **B**. Scale bars 40 µm. **B.** Individual channels depicting the morphological alterations at indicated time points for astrocytes (**left**) and oligodendrocytes (**right**). Position of magnified cells marked with the dashed white box: The astrocyte died within the first hour of the experiment (50-60 min). In contrast, the oligodendrocyte soma swelled, and its processes beaded, followed by the appearance of a prominent nucleus (at 280 min and 430 min, prominent nucleus, PN; pink arrowheads). Only a subset of the swollen oligodendrocytes is lost at a later time (at 360 min and 440 min, gray arrowheads). Scale bars 30 µm. **C.** Survival plot of glial cells following AQP4-IgG and complement application for 8 hours. Oligodendrocyte injury develops slowly (purple) compared to swift astrocyte depletion (green). Mean ± SEM (441 astrocytes, 201 oligodendrocytes; N = 3 mice). **D. Top:** Line diagram showing the time course of the onset of prominent nuclei (Olig_PN_) and cell loss (Olig_Loss_) for the subset of oligodendrocytes that died during 8-hour time-lapse recordings. Cells are ordered by time of onset of the Olig_PN_ state. **Bottom:** Dot-plot showing the time from the Olig_PN_ to Olig_Loss_. Mean ± SEM (37 oligodendrocytes; N = 3 mice).

### Oligodendrocyte injury in NMOSD is distinct and independent from astrocyte pathology

To understand the cellular injury mechanisms and the potential role of astrocyte loss in oligodendrocyte pathology, we investigated the impact of antibody-independent astrocyte loss on the oligodendrocytes. We used two-photon laser ablation, which disrupts membranes (Brill *et al*., 2011; Snaidero *et al*., 2020), to kill a small cluster of astrocytes. Subsequently, oligodendrocytes were imaged for up to 6 hours, a time window during which, in experimental NMOSD lesions, oligodendrocyte pathology is clearly manifest. No signs of oligodendrocyte injury were observed in the region of astrocyte ablation (oligodendrocyte survival at 360 min = 100.0 ± 0%; **Fig. 2A-B**). Furthermore, to model a global astrocyte depletion comparable to levels observed in our NMOSD lesions, we employed a selective, toxin-based approach for targeted cell ablation as previously described (Feng *et al*., 2016). Intermedilysin (ILY) is a pore-forming toxin that selectively binds the human complement protein CD59 (huCD9), leading to the formation of cytolytic pores, while exhibiting no reactivity toward mouse CD59 (Hu *et al*., 2008; Nagamune *et al*., 1996). Using a floxed knock-in allele (Feng *et al*., 2016), we selectively overexpressed huCD59 in astrocytes. Local application of ILY for 1 hour resulted in widespread astrocyte loss. At the same time, oligodendrocyte somata were largely preserved within the same region (**Supp. Fig. 1**). Collectively, these findings indicate that mechanisms beyond the loss of local astrocytic support and their lytic destruction are involved in the spread of glial pathology.

**Figure 2.**
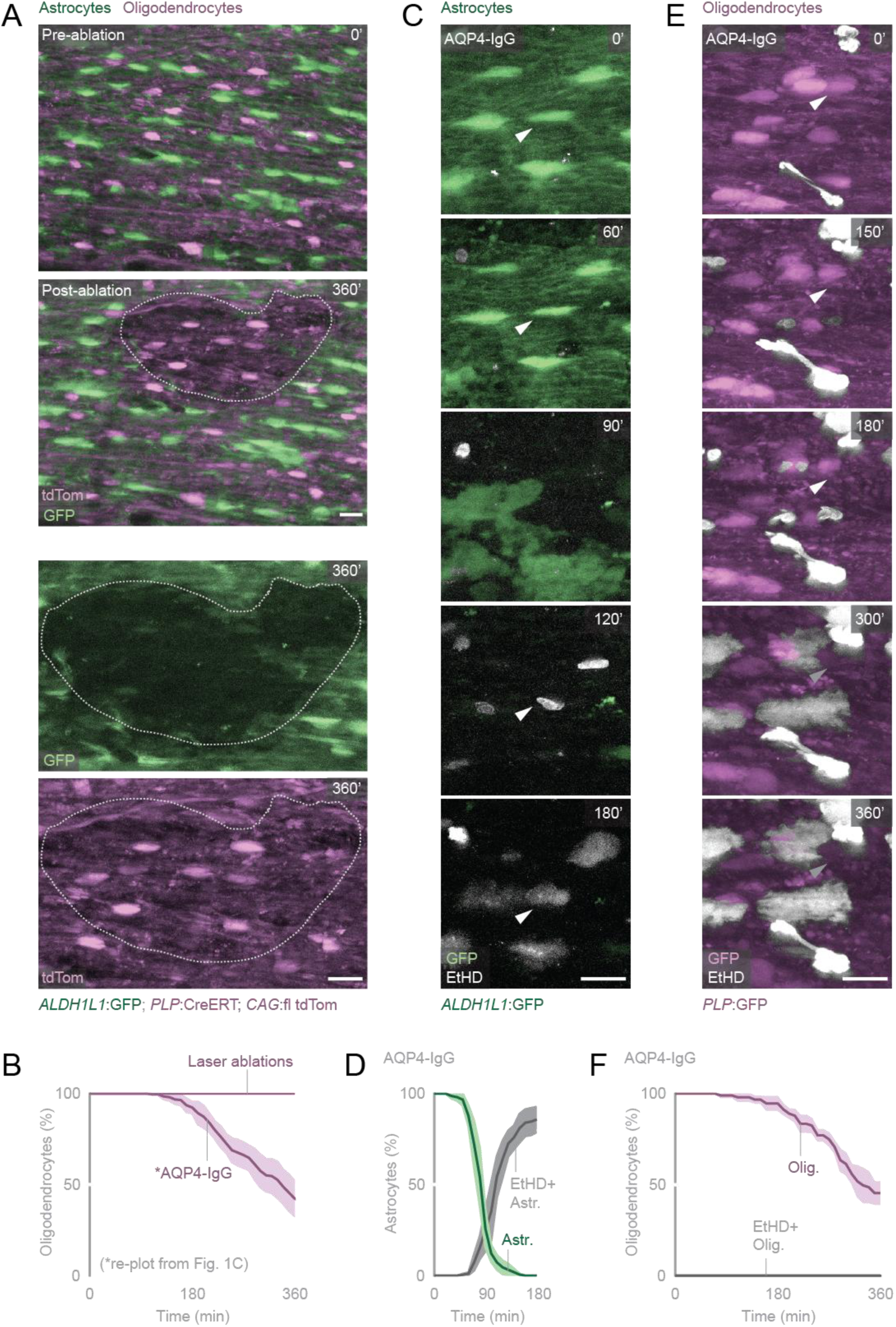
Oligodendrocyte injury is independent of and distinct from astrocyte damage. **A. Top:** Two-photon laser ablation of astrocytes (green) does not lead to morphological alteration or loss of oligodendrocytes (magenta). **Bottom:** Magnified view of the individual channels for astrocytes (GFP) and oligodendrocytes (tdTom) at 360 min after laser ablation. Scale bars 20 µm. **B.** Quantification of oligodendrocyte survival 6 hours following either AQP4-IgG and complement application or laser ablation of astrocytes. Mean ± SEM (44 oligodendrocytes, N=3 mice; AQP4-IgG line is replotted from Fig. 1C). **C.** Astrocyte death observed via the loss of cytoplasmic GFP signal induced by AQP4-IgG/complement application. Astrocytes (arrowhead) take up the cell-impermeable nuclear dye, ethidium homodimer (EtHD; gray). Scale bar 20 µm. **D.** Quantification of astrocyte survival (green) and ethidium homodimer uptake (gray). Mean ± SEM (149 astrocytes; N = 4 mice). **E.** Oligodendrocyte injury in AQP4-IgG/complement-induced lesions. Oligodendrocytes are not labeled with EtHD in contrast to the nuclei of other cells in the vicinity. Gray arrowheads indicate oligodendrocyte nuclei that remain EtHD negative even after the cytoplasmic GFP signal is lost and other cells (presumably astrocytes) have been labeled. Cells labeled with EtHD at 0 minutes are likely due to dura removal. Scale bar 20 µm. **F.** Quantification of oligodendrocyte survival (purple) and nuclear EtHD uptake in these cells (gray) – which runs as a “zero” line across the x-axis. Mean ± SEM (93 oligodendrocytes; N = 3 mice).

To further compare injury mechanisms and identify a divergence point in cell death pathways, we assessed the extent of membrane disruption during AQP4-IgG/complement exposure. As shown before (Brill *et al*., 2011; Herwerth *et al*., 2016), ethidium homodimer (EtHD), a membrane-impermeable nuclear dye, readily labeled the nuclei of astrocytes after AQP4-IgG mediated injury, once GFP was lost (*ALDH1L1*:GFP; astrocyte loss t_50%_ = 78.6 ± 5.3 min; EtHD^+^ astrocytes t_50%_ = 109.0 ± 8.1; **Fig. 2C-D**). In contrast, oligodendrocytes (*PLP*:GFP; Mallon *et al*., 2002) did not take up EtHD at 360 min (oligodendrocyte survival = 45.5 ± 6.1%; EtHD^+^ oligos = 0.0 ± 0.0%; **Fig. 2E-F**). This suggests that membrane integrity, at least in the peri-nuclear region, was relatively preserved. Together, these data point to an oligodendrocyte injury mechanism that extends beyond simple astrocyte loss and involves variable degrees of membrane disruption.

### Oligodendrocytes exhibit calcium dyshomeostasis early in experimental NMOSD lesions

Putative mechanisms of oligodendrocyte injury involve glutamate excitotoxicity (Wrzos *et al*., 2014), calcium-mediated myelin vesiculation via unresolved toxic spill-over factors (Weil *et al*., 2016), and complement-mediated “bystander” membrane poration (Tradtrantip *et al*., 2017). All these processes are expected to involve changes in calcium signaling in oligodendrocytes. Therefore, we assayed glial calcium dynamics to determine whether intracellular calcium signals are related to glial pathology in experimental NMOSD. Cell-specific expression of a genetically encoded calcium sensor, GCaMP5g (floxed reporter mouse line GCaMP5G-IRES-tdTomato; Akerboom *et al*., 2012; Gee *et al*., 2014) in oligodendrocytes (*PLP*:CreERT; Doerflinger *et al*., 2003) or astrocytes (*GFAP*:Cre; Gregorian *et al*., 2009) allowed monitoring intracellular calcium in these two glial populations. In these measurements, we refrained from using NMOSD plasma as an antibody source and instead used a set of recombinant antibodies (recombinant AQP4 antibody, rAQP4-IgG, clone 7-5-53; or isotype control antibody r-Ctr-IgG, clone ICOS-5-2; Bennett *et al*., 2009) to minimize the interference of other biological components present in plasma samples that could elicit calcium signaling. In previous experiments, we established the equivalence of astrocytic lesions induced by NMOSD plasma and recombinant IgG (Herwerth *et al*., 2016; Herwerth *et al*., 2022). Both astrocytes and oligodendrocytes showed an intracellular calcium rise soon after the application of rAQP4-IgG with complement, even before morphological alterations became apparent in either glial population (**Fig. 3**). In astrocytes, the calcium rise was rapid and global (astrocytic high Ca^2+^ t_50%_ = 39.5 ± 7.4 min; **Fig. 3A-B**; **Supp. Video 3**), observed before lytic demise (t_50%_ = 78.1 ± 16.8 min). Control r-Ctrl-IgG treatment induced no changes in astrocytic morphology or calcium levels (**Fig. 3A-B**). Oligodendrocytes, in contrast, showed local and transient calcium elevations, often in processes, but also in the soma (**Fig. 3C**, **3E**; **Supp. Video 4**). These local signals then progressed to a global, steady-state high-calcium condition across the entire cell (at least in three consecutive imaging frames during the time-lapse, t_50%_ = 58.5 ±19.2 min). This lasting and pan-cellular high-calcium state typically preceded the granular transformation of oligodendrocyte processes and the appearance of a prominent nucleus in the soma (**Fig. 3C).** As for astrocytes, r-Ctrl-IgG and complement did not induce morphological changes or substantial steady state alterations in calcium signaling. Notably, in some control experiments, a minor population of oligodendrocytes still showed signs of treatment-induced calcium signals. This may indicate the presence of signaling-active ligands in the plasma-derived complement preparation or a special sensitivity of oligodendrocytes to low levels of complement activation even under control conditions (Liu *et al*., 2016). Despite this background signal, our experiments show that while astrocytes are the primary target of complement-mediated membrane disruptions, oligodendrocytes also exhibit changes in intracellular calcium signaling. Such signals would either be compatible with a specific signaling event induced by astrocyte loss or with sublytic membrane disruptions that allow calcium influx but do not progress to the overt membrane disruptions detected by EtHD.

**Figure 3.**
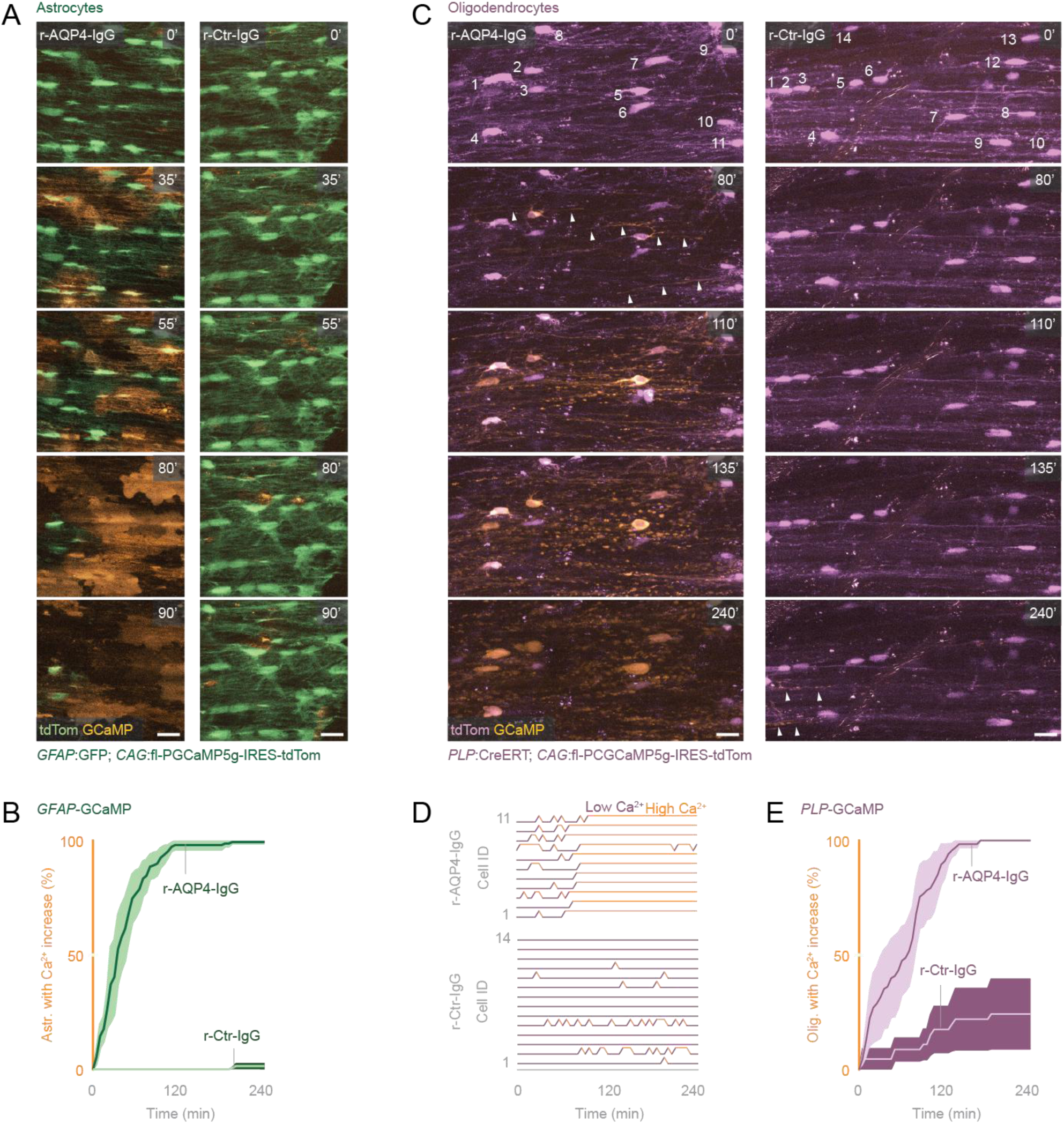
Glial cells show distinct calcium dynamics after NMOSD-related injury. **A.** In vivo time-lapse imaging of astrocytes (green) expressing the calcium sensor, GCaMP5 (orange), shows global and persistent intracellular calcium elevations following rAQP4-IgG and complement treatment. In control experiments, few astrocytes have a stable calcium rise, with sparse calcium transients restricted to astrocyte processes. Scale bars 20 µm. **B.** Percentage of astrocytes with persistently high calcium levels (>1.5x to baseline GCaMP signal, measured in the astrocyte somata) following application of rAQP4-IgG or r-Ctr-IgG with complement. Mean ± SEM (AQP4-IgG, 157 astrocytes; N=4 mice; r-Ctr-IgG, 123 astrocytes; N = 3 mice). **C.** GCaMP5 (orange)-expressing oligodendrocytes (magenta) show intracellular calcium uptake following rAQP4-IgG and complement treatment, which start as calcium transients in the oligodendrocyte processes (arrowheads at 80 min, not included in **D** and **E**). These preceded somatic calcium fluctuations that progressed to sustained high-calcium states, as well as morphological alterations, such as process beading (110 min) and prominent nuclei (135 min). Under control conditions, calcium signaling in processes and somata is mostly transient. Numbering of oligodendrocyte soma refers to plots in **D**. Scale bars 20 µm. **D.** Binarized calcium states (scored as high >1.5x to baseline, orange; otherwise, low, purple) for oligodendrocyte somata in **C** (n = 11 cells in rAQP4-IgG; n = 14 cells in r-Ctr-IgG). While somatic calcium transients also occur occasionally under control conditions, persistent calcium states are restricted to the rAQP4-IgG group. **E.** Percentage of oligodendrocytes with steady state high calcium levels in the soma (>1.5x to baseline, at least in three consecutive time points) following 4-hour rAQP4-IgG or r-Ctr-IgG and complement application. Mean ± SEM (rAQP4-IgG, 69 oligodendrocytes; N = 5 mice; r-Ctr-IgG, 46 oligodendrocytes; N = 4 mice).

### The MAC-inhibitor huCD59 protects oligodendrocytes from AQP4-IgG-mediated injury

To directly test the possibility of sublytic membrane disruptions, which can occur when activated complement proteins spill over from the primary target of an autoantibody attack to another (Park *et al*., 1997), we employed a genetic system that provides cell-type-specific and cell-autonomous protection against complement-mediated lysis. AQP4-IgG-mediated complement targeting of astrocytes involves the enzymatic activation of the complement cascade, culminating in the sequential insertion of soluble C5b to C9 proteins into the plasma membrane, forming an osmolytic pore known as the membrane attack complex (MAC, C5b-9). These pores cause rapid necrotic lysis by allowing ion and water flux across the membrane of antibody-targeted cells (Bayly-Jones *et al*., 2017). However, the soluble portion of the activated complement proteins can also lead to off-site targeting, i.e., “bystander injury”, which, based on in vitro studies and tissue stainings, has been suggested to contribute to axon-glial injury in NMOSD (Duan *et al*., 2018; Tradtrantip *et al*., 2017). While our previous analysis of axon injury in our model has demonstrated preserved membrane integrity and hence ruled out pore formation in axons (Herwerth *et al*., 2022), the possibility of off-target bystander injury as an in vivo mechanism of oligodendrocyte pathology in NMOSD has not been thoroughly investigated. To achieve this, we established a system of cell-type-specific transgenic overexpression of the human membrane attack complex (MAC) inhibitor CD59 (Feng *et al*., 2016). CD59 is a glycosylphosphatidylinositol-anchored membrane protein that blocks MAC pore formation by interfering with the oligomerization and incorporation of C9 protein into the nascent complex (Farkas *et al*., 2002; Kimberley *et al*., 2007). Additionally, previous studies have documented that, in mice, overexpression of huCD59 in endothelial cells and in some hematopoietic cells, such as platelets, neutrophils, and monocytes, protects against MAC-mediated atherosclerosis and aneurysms (Wu *et al*., 2010; Wu *et al*., 2009).

First, to establish the inhibitory function of huCD59 overexpression on antibody-mediated MAC-driven injury in our system, we performed AQP4-IgG/complement experiments with huCD59 overexpression on astrocytes (*GFAP*:Cre x *CAG*:fl tdTom x *CAG*:fl huCD59). Astrocytes expressing huCD59, even in heterozygous animals for the huCD59 allele, were well protected against direct complement attack up to 6 hours of lesion induction (astrocyte survival%: huCD59_HE_ = 90.0 ± 3.3; huCD59_HO_ = 92.6 ± 3.0;; **Fig. 4A-C**). The protection of astrocytes also prevented secondary oligodendrocyte injury in these animals (astrocyte survival%: huCD59_HE_ = 87.5 ± 5.3; oligodendrocyte survival%: 100.0 ± 0.0; **Fig. 4D-E**). In contrast, huCD59 negative littermate controls showed complete astrocyte depletion after 3 hours (WT = 1.4 ± 0.9%; **Fig. 4A-C**), similar to the initial experiments we performed in *ALDH1L1*:GFP; *PLP*:CreERT x *CAG*:fl tdTom mice (**Fig. 1**).

**Figure 4.**
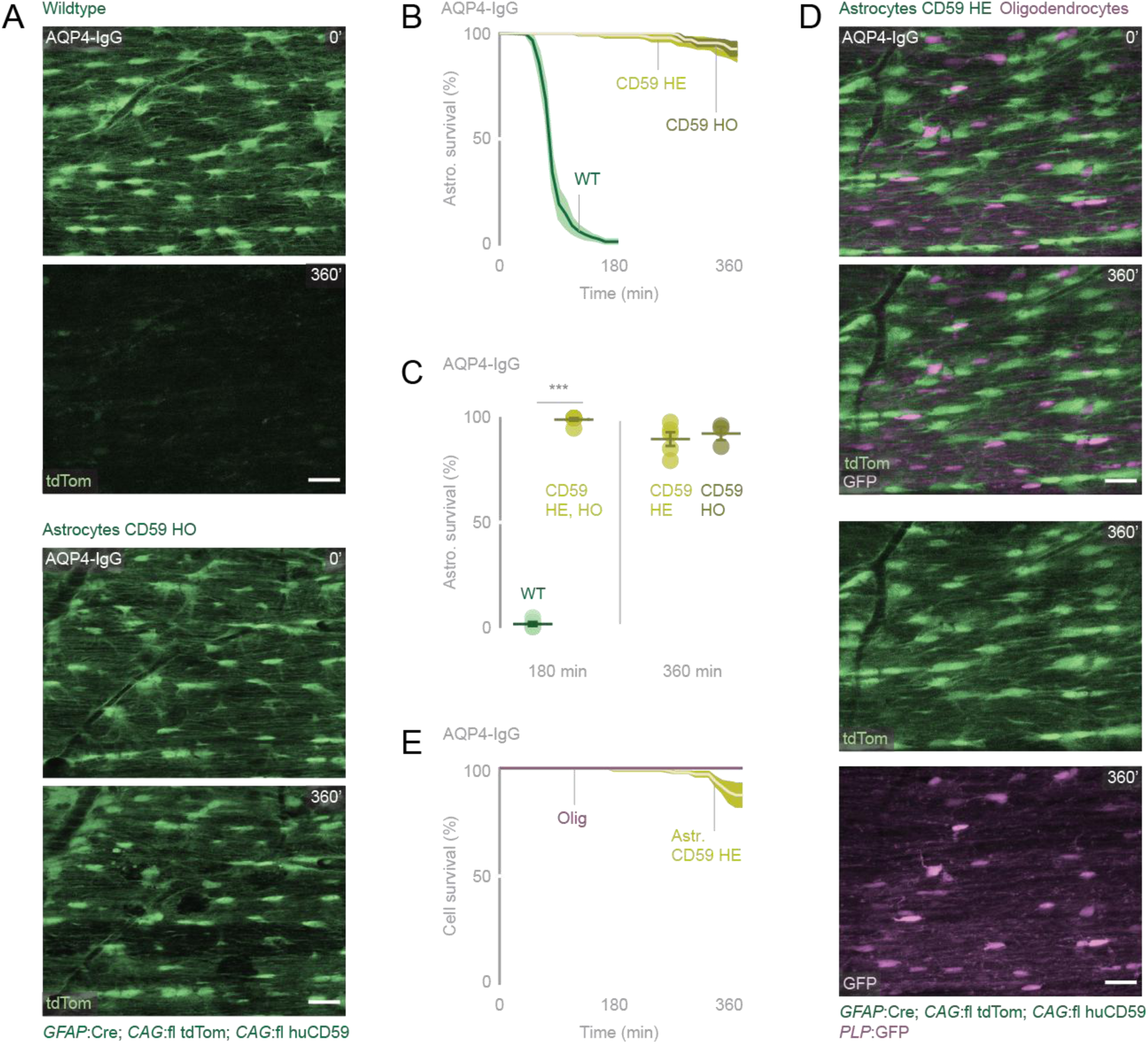
Astrocytic expression of the MAC inhibitor huCD59 protects against AQP4-IgG-mediated injury. **A.** Following 6-hour treatment with AQP4-IgG and complement, astrocytes expressing huCD59 were protected from complement-mediated attack and cell lysis in GFAP:Cre; CAG:fl tdTom; CAG:fl huCD59 mice (CD59 homozygous, HO). Astrocytes lacking huCD59 expression were largely depleted after 3 hours of AQP4-IgG/complement exposure in littermate controls (huCD59 negative mice, wildtype). Scale bars 30 µm. **B.** Astrocyte survival over time. A high astrocyte survival rate was observed in huCD59 homozygous (140 astrocytes, N=3 mice) and heterozygous mice (242 astrocytes; N = 5 mice) compared to wildtype littermate controls (238 astrocytes; N = 5 mice). **C.** Fraction of surviving astrocytes at 3-hour and 6-hour. huCD59 negative littermate controls showed almost complete astrocyte loss (N = 5 mice). Pooled data from CD59 HO and HE mice showed robust astrocyte survival at 3-hour. Data are shown as mean ± SEM. Mann–Whitney test: ***p = 0.0008 (WT vs pooled CD59 HE/HO at 180 min). **D.** Representative in vivo time-lapse images demonstrating the protection of glial cells, astrocytes (green) and oligodendrocytes (magenta) from AQP4-IgG-mediated injury in *GFAP*:Cre; *CAG*:fl tdTom; *CAG*:fl huCD59; *PLP*:GFP mice. Scale bars 30 µm. **E.** Survival of oligodendrocytes (*PLP*:GFP; magenta) and huCD59-expressing astrocytes (yellow) following 6-hour of AQP4-IgG/complement treatment in *GFAP*:Cre; *CAG*:fl tdTom; *CAG*:fl huCD59; *PLP*:GFP mice. The protection of astrocytes also prevented secondary oligodendrocyte injury in these animals (157 astrocytes, 115 oligodendrocytes; N = 3 mice).

Next, to test the off-target effects of the complement, huCD59 was selectively overexpressed in oligodendrocytes (*MOG*:Cre, Hovelmeyer *et al*., 2005, crossed with *CAG*:fl tdTom x *CAG*:fl huCD59). Strikingly, oligodendrocyte injury following AQP4-IgG/complement was completely blocked (oligodendrocyte survival%: WT = 26.6 ± 2.2; huCD59_HE_ = 96.1 ± 0.9; huCD59_HO_ = 100.0 ± 0.0; **Fig. 5A-C**). To ensure that the depletion of astrocytes was not hindered in these experiments, quadruple transgenic mice with dual labelling of both the astro- and oligodendroglial cell population were used (*ALDH1L1*:GFP x *MOG*:Cre x *CAG*:fl tdTom x *CAG*:fl huCD59; **Fig. 5D-E**). We observed no changes in the degree of astrocyte depletion after 6 hours (1.2 ± 0.8%, **Fig. 5E**). Overall, our results provide an unambiguous *in vivo* demonstration of off-target complement targeting of acute NMOSD-related oligodendrocyte pathology.

**Figure 5.**
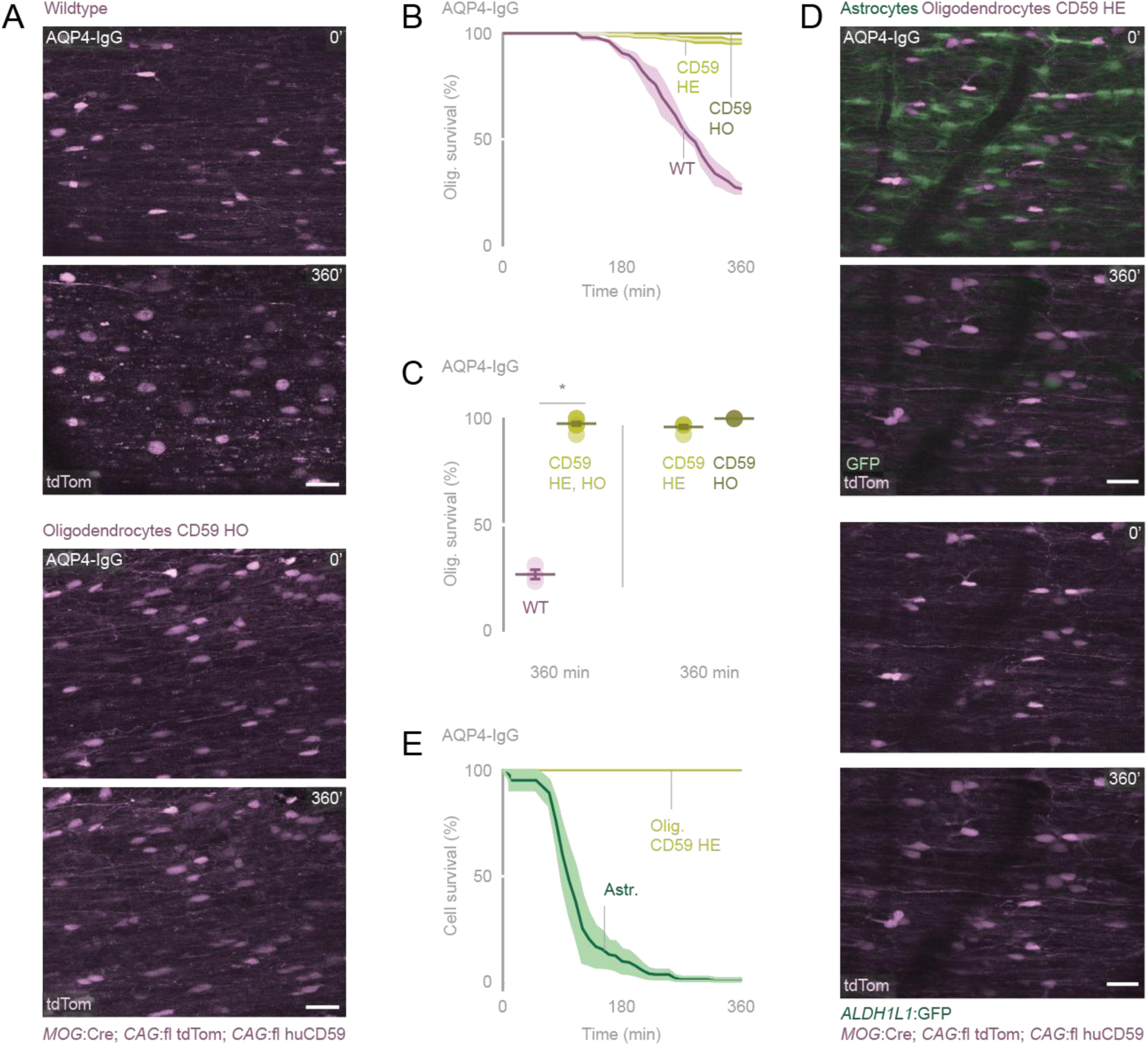
MAC inhibitor huCD59 expression protects oligodendrocytes despite AQP4-IgG–mediated astrocyte injury. **A.** Following 6-hour treatment with AQP4-IgG and complement, minimal injury was observed in oligodendrocytes overexpressing huCD9 (MOG:Cre; CAG:fl tdTom; CAG:fl huCD59 mice; CD59 HE and HO mice). Scale bars 30 µm. **B.** Oligodendrocyte survival over time. A high survival rate was observed in huCD59 homozygous (129 oligodendrocytes; N = 3 mice) and heterozygous mice (176 oligodendrocytes; N = 5 mice) compared to wildtype littermate controls (104 oligodendrocytes; N = 3 mice). **C.** Fraction of surviving oligodendrocytes at 6-hour. CD59 HO and HE mice showed robust survival at 6-hour compared to huCD59 negative littermate controls with substantial oligodendrocyte injury (N = 3 mice). Data are shown as mean ± SEM. Mann–Whitney test: *p = 0.0121 (WT vs pooled CD59 HE/HO at 360 min). **D.** Representative in vivo time-lapse images demonstrating the protection of oligodendrocytes (magenta) despite AQP4-IgG-mediated astrocyte depletion in *MOG*:Cre; *CAG*:fl tdTom; *CAG*:fl huCD59; *ALDH1L1*:GFP mice. Scale bars 30 µm. **E.** Survival of astrocytes (*ALDH1L1*:GFP; green) and huCD59-expressing oligodendrocytes (CD59 HE, yellow) following 6 hours of AQP4-IgG/complement treatment in MOG:Cre; CAG:fl tdTom; CAG:fl huCD59; ALDH1L1:GFP mice (164 astrocytes, 107 oligodendrocytes; N = 4 mice).

### Off-target complement injury is not a general phenomenon among other glial cells

While the off-target complement activation clearly induces oligodendrocyte injury, it remains unclear whether this represents generalized toxicity affecting all glial cells or a spill-over phenomenon specific to oligodendrocytes. To address this, we investigated the responses of additional glial cell types within acute lesions, specifically microglia (*CX3CR1:*GFP knock-in mice; Jung *et al*., 2000) and oligodendrocyte precursor cells (OPCs; *CSPG4:*DsRed transgenic mice; Zhu *et al*., 2008) during 6 hours of AQP4-IgG/complement treatment. Microglia rapidly became activated, extending processes and forming protrusions for engulfment, which likely contributed to debris removal following astrocyte lysis (100.0 ± 0; **Fig. 6A**; Vidal-Itriago *et al*., 2022) – but no cell loss was observed. Similarly, the OPC population was largely unaffected within the lesion site, with only a minor reduction in cell number (OPC survival at 360 min: rAQP4 = 91.1 ± 4.5; ctr-IgG = 100.0 ± 0.0; **Fig. 6B-C**). The *CSPG4:*DsRed transgenic line also labels perivascular cells, likely pericytes, which can be distinguished from OPCs by their morphology and localization (**Fig. 6B**). Pericyte loss was observed in both the AQP4-IgG and control treatment groups to varying degrees, likely reflecting the sensitivities arising from dura removal (pericyte survival%: rAQP4 = 61.4 ± 8.1; ctr-IgG = 48.3 ±19.5). Together, these findings indicate that the effects of off-target complement-mediated injury within the glial population are largely restricted to oligodendrocytes. Together with our previous finding that axons do not undergo early membrane poration in experimental NMOSD lesions (Herwerth *et al*., 2022), this suggests a high degree of cellular specificity of bystander injury, despite its origin in the “non-specific” off-target effects of the cell-type-restricted antibody-complement attack. This raises the question of whether this phenomenon originates in an especially vulnerable cellular relationship that astrocytes and oligodendrocytes share in vivo.

**Figure 6.**
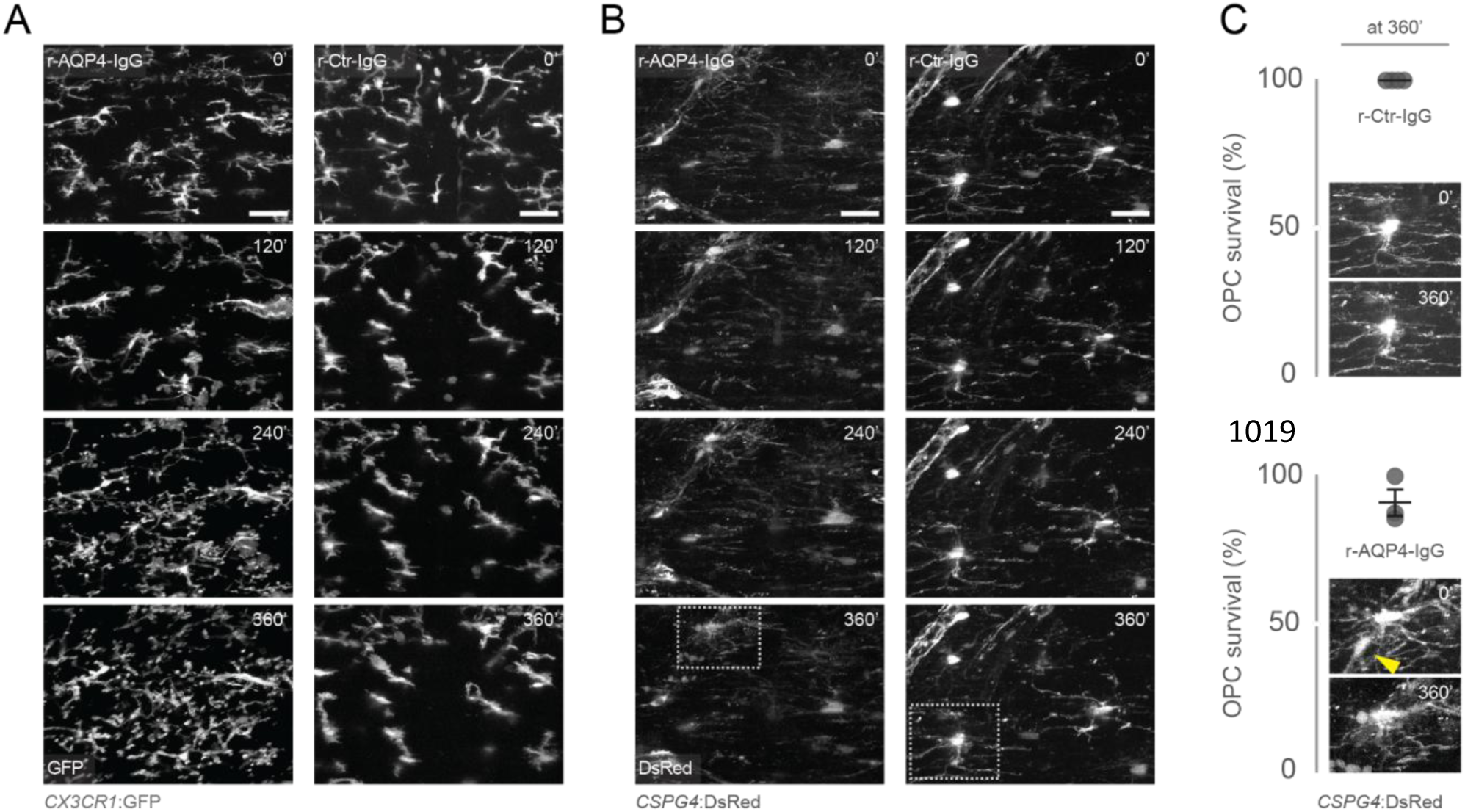
Microglia and oligodendrocyte precursor cells are spared in experimental NMOSD lesions. **A.** Individual time frames showing microglial activation (*CX3CR1*:GFP) in the presence/context of AQP4-IgG and complement, characterized by process extension and the formation of engulfment protrusions (**left**). In control experiments, microglia maintained their morphology without process extension (**right**). Scale bars 30 µm. **B.** Oligodendrocyte precursor cells (OPCs; *CSPG4*:DsRed) remain mostly unaffected in both AQP4-IgG (left) or control conditions (**right**). Dashed white boxes indicate the cells shown in panel C. **C.** Fraction of surviving OPCs at 6-hour following r-Ctr-IgG (N = 4 mice) or rAQP4-IgG (N = 3 mice) treatment. Representative OPCs from each experiment are shown in the insets. The yellow arrowhead indicates a pericyte along the vasculature, identifiable by its elongated cell body, in contrast to the characteristically round soma of OPCs. Data are shown as mean ± SEM.

### Complement injury spreads bidirectionally between astrocytes and oligodendrocytes

To investigate this potential sensitivity of the astrocyte-oligodendrocyte pairing to complement-mediated attack, another antibody-based approach was used to primarily target oligodendrocytes. For this, we used a MOG-IgG/complement combination (humanized recombinant MOG antibody, rMOG-IgG 8-18C5 (Beltran *et al*., 2021; Brandle *et al*., 2016; Linington *et al*., 1988), which targets a myelin and oligodendrocyte surface antigen, myelin oligodendrocyte glycoprotein (MOG), and can, in its murine form, mediate complement-mediated demyelination in rodents (Linington *et al*., 1988; Romanelli *et al*., 2016). We then imaged both astrocytes and oligodendrocytes in parallel for up to 6 hours (*ALDH1L1*:GFP; *PLP*:CreERT x *CAG*:fl tdTom mice; **Fig. 7**). This resulted not only in the expected oligodendrocyte injury (oligodendrocyte survival at 360 min: 13.3 ± 3.4%) but also in substantial astrocyte loss (astrocyte survival: 18.0 ± 10.7% **Fig. 7B**). The temporal profile of astrocyte loss (t_50_% = 266.0 ± 34.9 min; **Fig. 7B**) closely followed oligodendrocyte pathology, albeit with a small delay (t_50_% = 242.2 ± 19.0 min; **Fig. 7B**). Together, these findings support the notion that off-target complement injury can in principle be bidirectional between the major macroglial populations of the spinal cord.

**Figure 7.**
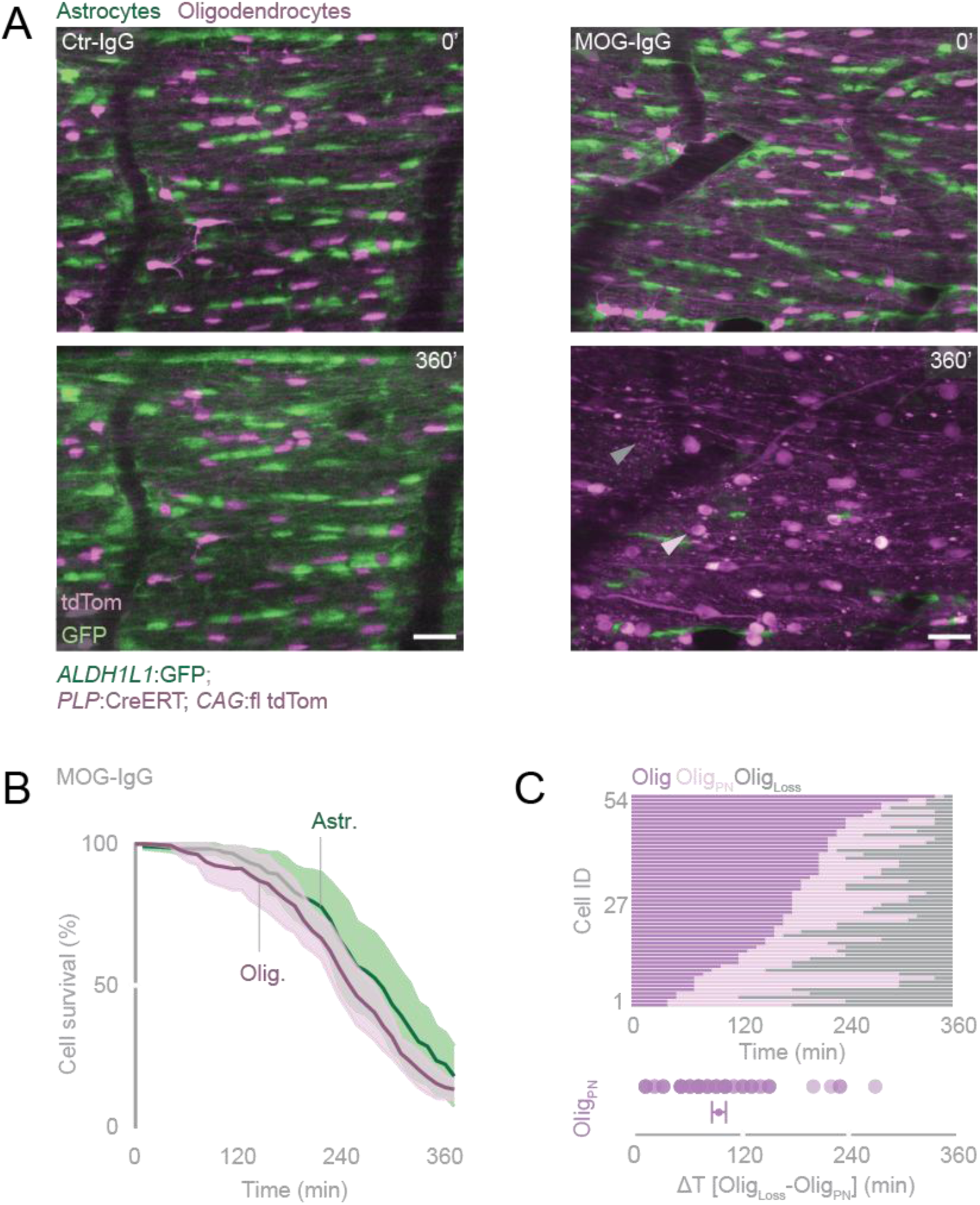
Astrocyte loss follows oligodendrocyte injury in MOG-IgG-mediated lesions. **A.** Spinal cord in vivo imaging of lesions targeting oligodendrocytes via MOG-IgG and complement (**right**; astrocytes, green; oligodendrocytes, magenta). Both astrocytes and oligodendrocytes are lost under these conditions. In control experiments (**left**), astrocytes and oligodendrocytes remain unaffected. Gray arrowhead indicates an oligodendrocyte lost during the 6-hour imaging period; pink arrowhead shows an oligodendrocyte displaying a prominent nucleus. Scale bars 30 µm. **B.** Survival plot of glial cells following MOG-IgG and complement application for 6 hours. Oligodendrocyte loss (purple) starts within the first hour, swiftly followed by astrocyte depletion (green). Mean ± SEM (195 astrocytes, 155 oligodendrocytes; N = 3 mice). **C. Top:** time course of the onset of prominent nuclei (Olig_PN_) and cell loss (Olig_Loss_) for the subset of oligodendrocytes that died during 6-hour time-lapse recordings. Individual data plots are ordered by the onset of the Olig_PN_ state. **Bottom:** Dot-plot showing the time from the Olig_PN_ to Olig_Loss_. Mean ± SEM (54 oligodendrocytes; N = 3 mice).

## Discussion

Early descriptions of NMOSD (under the eponym “Devic’s disease”; Lucchinetti *et al*., 2014) emphasized the demyelination that accompanies attacks in the optic nerve and spinal cord. While originally classified as a primary demyelinating disease, NMOSD was later redefined as a primary astrocytopathy following the discovery of AQP4-IgGs (Lennon *et al*., 2005; Lennon *et al*., 2004). Hence, attention shifted to astrocytes and their AQP4-enriched processes as the initial site of pathology. In contrast, oligodendrocyte injury was often attributed to either immune-mediated damage or chronic glial dysfunction resulting from the loss of astrocytic homeostatic support (Lucchinetti *et al*., 2014). Yet, this notion of purely “secondary” oligodendrocyte damage is challenged by the rapid onset and the extent of demyelination observed in acute lesions, where oligodendrocyte injury can emerge within hours (Wrzos *et al*., 2014). By directly imaging lesion formation and pathology spread in an experimental NMOSD lesion model, we now resolve this discrepancy: AQP4-IgG-mediated complement activation does not remain locally confined to astrocytes but initiates a rapid second wave of an off-target complement injury that reaches oligodendrocytes. These findings provide a mechanistic explanation for the strong demyelinating component of NMOSD, while also conceptually reconciling the histopathological picture (and early descriptions) of the disease with its current pathomechanistic interpretation.

A striking feature of the murine spinal cord lesions is the rapid damage of oligodendrocytes following AQP4-IgG/complement-mediated injury to astrocytes. With the initial attack clearly directed at astrocytes (Herwerth *et al*., 2016), oligodendrocytes exhibit concomitant fluctuations in intracellular calcium levels soon after the initial signs of astrocyte injury observed (t_50_[Olig_High Ca2+_]= 58.5 ±19.2 min; **Fig. 3E**). Within hours of astrocyte lysis, oligodendrocytes undergo structural alterations and subsequently often die (t_50_[Olig_survival_] = 340.6 ± 29.6 min; **Fig. 1C, D**) in a timeline that happens before immune cells invade in similar experimental lesions (typically at >8h; Wrzos *et al*., 2014). While this tight temporal coupling makes it unlikely that secondary immunological effects in evolving lesions are the initial cause of demyelination, several explanations remain compatible with the development of oligodendrocyte injury following astrocytic loss (Lucchinetti *et al*., 2014). This includes mechanisms such as impaired water and ion buffering to which oligodendrocytes are vulnerable (Nicaise *et al*., 2019), gap junction-mediated calcium transfer, as astrocytes and oligodendrocytes form a network via these conduits (Masaki, 2015), or injury-related glutamate excitotoxicity (Marignier *et al*., 2010; Wrzos *et al*., 2014). Arguing against such mechanisms, our in vivo laser ablation of astrocytes, which is expected to cause membrane disruptions and lytic cell death (Breckwoldt *et al*., 2014; Brill *et al*., 2011), leaves oligodendrocytes unaffected. Similarly, ILY-toxin-mediated astrocyte poration was also largely innocuous for oligodendrocytes (**Supp. Fig. 1**). This rules out the possibility that acute oligodendrocyte damage arises simply from the loss of astrocyte function or the release of toxic factors in our NMOSD model. This is despite substantial microglial activation, which is a common response to CNS laser lesions (Nimmerjahn *et al*., 2005) and is also a prominent feature of our AQP4-IgG/complement induced lesions (**Fig. 6A**). Thus, our results suggest another form of acute, secondary damage to oligodendrocytes, which, however, exhibits different temporal characteristics and is not overtly necrotic, given its relatively slow manifestation and the subtle morphological alterations when compared to astrocytes. At the same time, the initiating event is clearly the direct engagement of complement (specifically the MAC) with oligodendrocyte membranes, as demonstrated by the striking protective effect of cell-type-specific expression of human CD59, a GPI-anchored inhibitor protein that blocks MAC assembly (Farkas *et al*., 2002; Kimberley *et al*., 2007). When expressed only in oligodendrocytes, huCD59 prevented oligodendrocyte injury despite the substantial loss of astrocytes (**Fig. 5D-E**). Moreover, in line with previous work on systemic models of NMOSD (Wang *et al*., 2017), expressing huCD59 in astrocytes preserved both astrocytes and neighboring oligodendrocytes (**Fig. 4D-E**), indicating that complement activation must occur first at the AQP4/AQP4-IgG complex on the astrocyte membrane; therefore, ruling out direct engagement of the AQP4-IgG with oligodendrocyte membranes. Together, these experiments reveal a spatially and temporally ordered chain of events: complement activation is initiated at astrocytes but not restricted to them; instead, soluble complement intermediates or sublytic MAC complexes propagate to adjacent oligodendrocytes, causing functional injury while sparing other nearby cell types.

Notably, such off-target or “bystander” phenomena have been observed in vitro in cellular models of NMOSD-related injury (Duan *et al*., 2018; Tradtrantip *et al*., 2017). Furthermore, the presence of MAC on oligodendrocytes has been suggested based on light microscopy analysis, while the upstream complement proteins in the cascade were only present on astrocytes (Tradtrantip *et al*., 2017), although diffraction-limited resolution may be insufficient to pinpoint the exact cellular location of such molecular assemblies in the crowded environment of the neuropil (Mezydlo *et al*., 2023). Thus, it appears that substantial complement deposition can compromise the precise targeting and effector functions of the complement system in the CNS, leading to bystander damage to oligodendrocytes. At the same time, this form of “off-target” injury exhibits a degree of specificity. Microglia and oligodendrocyte precursor cells, despite their proximity to the injury site, remain structurally and functionally intact. Microglia were not injured but rather activated (**Fig. 6A**). Moreover, only a minor population of OPCs showed minimal signs of damage (**Fig. 6C**, <10%), which could be explained by rare OPC differentiation patterns with AQP4 expression as previously described in neuroinflammatory conditions in both experimental and human datasets (Blaszczyk *et al*., 2025; Jakel *et al*., 2019) or early myelinating transition states that already exhibit features that render them vulnerable to MAC spillover from nearby astrocytes. This hierarchy of injury across glial cell types recapitulates what is seen in human NMOSD lesions (Wrzos *et al*., 2014) and suggests that complement-mediated propagation is not a simple diffusion-driven process but rather interacts with the intrinsic features of mature oligodendrocytes. This mechanism of glial propagation also illuminates further aspects of the architecture observed in early NMOSD lesions. Regions with high AQP4 density, such as perivascular endfeet, sustain intense complement activation and rapid astrocyte lysis. Surrounding myelinated segments, by contrast, may be exposed to sublytic complement complexes that trigger oligodendrocyte dysfunction without immediate cell death. This establishes a lesion topology in which astrocytic and oligodendrocytic injury zones overlap but are not identical, and in which secondary demyelination spreads outward from the astrocytic core. Overall, the picture of a special relationship between astrocytes and oligodendrocytes emerges that might relate to the proximity and intertwining of their processes, e.g., near nodes of Ranvier (Marignier *et al*., 2010), where direct interactions have been described during myelin remodeling (Dutta *et al*., 2018; Lucchinetti *et al*., 2014). Indeed, such perinodal astrocyte processes, which in freeze-fracture electron micrographs contain orthogonal arrays of particles reminiscent of AQP4 (Black & Waxman, 1988) are the likely sites of off-target complement pathology.

Our work thus clarifies why NMOSD lesions consistently exhibit features of two distinct forms of glial death: Astrocytes show rapid calcium influx and swell quickly after MAC formation, consistent with their high local load of complement activation and membrane poration. Oligodendrocytes, in contrast, undergo a protracted decline characterized by sustained calcium elevation and nuclear condensation, and eventually die without overt lysis. These divergent dynamics of cellular demise despite a shared initial injury mechanism likely relate to a combination of differences in MAC pore load and intrinsic variabilities in complement defense capacity, calcium handling, or membrane repair mechanisms between the two glial types. Myelin membranes, with their unique composition, may alter the dynamics of MAC insertion (Couves *et al*., 2023), thereby delaying overt lysis of oligodendrocytes and rather promoting sublytic complement effects, which include calcium dyshomeostasis and activation of cell death pathways (Triantafilou *et al*., 2013). Similarly, different cell types exhibit differential levels of resilience against MAC insertion, including differential expression of various complement inhibitors (Dunkelberger & Song, 2010) and the ability to endocytose or shed membranes to remove MAC pores (Xie *et al*., 2020). Indeed, oligodendrocytes do not express substantial CD59 endogenously (Piddlesden & Morgan, 1993; Scolding *et al*., 1998; Wing *et al*., 1992). Furthermore, sublethal glutamate excitotoxicity - which may occur due to the almost complete depletion of astrocytes in our model - has been previously shown to mediate kainate receptor-driven calcium influx and ROS signaling, thereby sensitizing oligodendrocytes to complement-mediated attack (Alberdi *et al*., 2006; Matute, 2007). Thus, such differences in vulnerability and resilience could explain why oligodendrocyte injury is delayed, yet MAC assembly ultimately still results in myelin damage and cell death.

The presence of a prolonged sublethal phase in oligodendrocytes has therapeutic implications. During this period, oligodendrocytes retain structural integrity and could be rescued if complement activation is halted or calcium homeostasis is restored. Given that a single NMOSD attack can cause long-lasting disability (Berthele *et al*., 2023), timely intervention during the early complement-driven window may be critical for preserving oligodendrocytes and myelin. Moreover, a growing body of evidence suggests that under certain circumstances, damaged oligodendrocytes can contribute to remyelination, albeit to a debated extent (Bacmeister *et al*., 2020; Duncan *et al*., 2018; Mezydlo *et al*., 2023; Neely *et al*., 2022). This therapeutic window aligns with our prior observation of swift but potentially reversible axonal pathology in our NMOSD model (Herwerth *et al*., 2022) and in other forms of inflammatory and non-inflammatory spinal pathology (Williams *et al*., 2014; Witte *et al*., 2019).

A critical aspect of our findings is the bidirectionality of complement-driven propagation. When the initiating insult is shifted from astrocytes to oligodendrocytes using MOG-IgG, astrocytes become secondary victims of the complement attack. This reciprocity shows that a complement response can drive glial injury cascades in either direction, depending on which cell type is initially targeted. This insight relates to the clinico-pathological overlap and distinctions between NMOSD and MOGAD, two humoral autoimmune diseases that share early demyelinating features despite targeting distinct antigens. Our experimental results suggest that in both entities, the early mechanism of complement-mediated lesion expansion could, in principle, spread to the non-targeted, but spatially proximate, partner in the astrocyte-oligodendrocyte network. However, histopathological findings in MOGAD contradict this, showing that astrocyte loss is not prominent (Marignier *et al*., 2021). Also, pathology in MOGAD appears to be much less driven by complement and humoral factors in general. This might limit damage to myelin (which is exquisitely sensitive to calcium dyshomeostasis, as would be expected by sublytic membrane pores; Triantafilou *et al*., 2013), but typically does not cause overt loss of oligodendrocytes (or astrocytes; Hoftberger *et al*., 2020). In MOGAD, soluble GFAP – an astrocyte-enriched CSF biomarker – is elevated, albeit to a lesser extent than in NMOSD (Marignier *et al*., 2025), indicating the membranes of both glial cells might still be compromised. Further divergence, e.g., in immunological mechanisms or local glial networks, would be expected to contribute to the distinct chronic pathology and clinical outcome. For instance, in the cortex (which often demyelinates in MOGAD but rarely in NMOSD; Uzawa *et al*., 2024), astrocyte-oligodendrocyte contacts might be less compact than in classical white matter tracts, limiting the bystander mechanism we characterize here.

Several aspects of our model limit direct extrapolation to human inflammatory demyelinating conditions: we used a heterologous antibody-complement combination applied pially, whereas in patients, IgG and complement originate from the same species and usually enter the CNS via the vasculature, with occasional intrathecal production and pial access routes (Guo *et al*., 2017). Of note, MOG-IgG 8-18C5 used in our study binds to an epitope distinct from that targeted by human MOG-IgGs; therefore, the findings may not recapitulate human pathology (Peschl *et al*., 2017; *Saaodun et al.*, 2014). Moreover, secondary amplification processes (especially those related to the breakdown of the blood-brain barrier and the infiltration of immune cells; Winkler *et al*., 2021; Wrzos *et al*., 2014) are largely absent in our model. Indeed, our lesions lack the broader autoimmune context of NMOSD. In patients, antibody-mediated complement activation occurs alongside T-cell infiltration, granulocyte recruitment, and cytokine release, all of which may influence early glial vulnerability. Similarly, our model also does not elicit an endogenous complement response and hence cannot reflect all endogenous dynamics and regulation. Indeed, the evolution of experimental lesions in rodent NMOSD has been shown to depend on endogenous CD59 (Yao & Verkman, 2017a, b). Finally, differences between mouse and human spinal cord architecture mean that the spatial arrangements that enable complement spread between glial cells in our system may not be identical in human lesions. Thus, while our model reveals a mechanism that can operate in antibody-mediated CNS injury, it does not prove that all steps occur in this order or causality in human NMOSD lesions. Clarifying this applicability awaits further study in human tissue but is severely limited by the rarity of the disorder compared to other autoimmune CNS disorders, the decrease in biopsy indication given progress in biomarkers and clinical imaging, and consequently, the scarcity of acute lesion tissue, which is rarely obtained during autopsy.

Rather than directly addressing the human applicability of our model, we demonstrate the possibility of spread and the importance of considering such mechanisms of convergence when interpreting and targeting glial pathology. The fact that complement-mediated injury can spread almost seamlessly from targeted to non-targeted cells, with speed and specificity, supports a reframing of NMOSD as a primary gliopathy, in which the initiating autoimmune event is astrocyte-specific but the mechanism of lesion expansion rapidly encompasses oligodendrocytes. This does not diminish the role of AQP4-IgG in conferring disease specificity, but it emphasizes that the earliest pathological events could be shaped by complement biology rather than solely by astrocyte loss. Complement acts as the transglial conduit of injury, connecting antigen recognition with demyelination. Reframing a disease like NMOSD in this way has implications beyond taxonomy. It suggests that therapies should aim not only to block AQP4-IgG binding or systemic complement activation, but also to shield vulnerable glial populations during the brief hyperacute window when complement spreads.

## Materials and Methods

### Animals

Experiments were conducted using male and female mice aged 2-6 months. Animals were housed at 22 ± 2 °C with 50 ± 15% humidity under a 12-hour light/dark cycle. Astrocyte morphology was visualized using *ALDH1L1*:GFP (Yang *et al*., 2011; MGI:3843271) or *GFAP*:Cre (Gregorian *et al*., 2009; JAX: 24098) mice crossed with the Rosa26 *CAG*:STOP-flox-tdTomato reporter line (tdTom; Madisen *et al*., 2010; JAX: 7914). Oligodendrocyte morphology was assessed in *PLP*:GFP (Mallon *et al*., 2002; JAX: 33357) or *PLP*:CreERT-tdTom transgenic mice (*PLP*:CreERT; Doerflinger *et al*., 2003; JAX: 5975, crossed with tdTom; Madisen *et al*., 2010; JAX: 7914). For calcium imaging, *GFAP*:Cre and *PLP*:CreERT mice were crossed with Polr2a *CAG*:flox-GCaMP5g-IRES-tdTomato mice (GCaMP5g; Gee *et al*., 2014; JAX: 24477). Tamoxifen induction was not required due to constitutive Cre activity in *PLP*:CreERT progeny. Glia-specific expression of the human MAC inhibitor protein CD59 (huCD59) was achieved by crossing *MOG*:Cre-tdTom or *GFAP*:Cre-tdTom mice with Hipp11 *CAG*:STOP-flox-CD59 mice (huCD59; Feng *et al*., 2016, courtesy of Prof. Bin Gao, NIAAA, NIH, USA; *MOG*:Cre mice, Hovelmeyer *et al*., 2005, courtesy of Prof. Ari Waisman, University of Mainz, Germany). Microglia were imaged using *CX3CR1*:GFP knock-in mice (Jung *et al*., 2000, JAX: 5582). Oligodendrocyte precursor cells (OPCs) were imaged in *CSPG4*:DsRed transgenic mice (Zhu *et al*., 2008; JAX: 8241). All animal experiments were conducted in accordance with local regulations and approved by the relevant regulatory authorities.

### Complement and antibody sources

A plasma sample from an AQP4-IgG-positive NMOSD patient treated at the Klinikum rechts der Isar, Technical University of Munich, diagnosed according to established criteria (Wingerchuk *et al*., 2015), was used in this study. Control plasma and complement were obtained from healthy donors via the Bavarian Red Cross. Written informed consent was obtained from all participants. Plasma and serum samples were collected and processed as previously described (Herwerth *et al*., 2016; Herwerth *et al*., 2022). To eliminate complement activity, NMO- and control-plasma samples were heat-inactivated and used solely as IgG sources. Complement activity was supplied by pooled sera from three healthy donors. For AQP4-antibody experiments, a set of recombinant human IgG1 antibodies was used: rAQP4-IgG (rAb7-5-53), derived from a clonotypic plasmablast from NMO patient CSF (Bennett *et al*., 2009), and rCtrl-IgG (rAb ICOS-5-2), an IgG1 of unknown specificity from a chronic meningitis patient, served as an isotype control. For MOG antibody experiments, a humanized recombinant derivative of the mouse monoclonal antibody 8-18C5 (IgG1; MOG-IgG), incorporating human IgG1 and κ constant regions, was used (Beltran *et al*., 2021; Brandle *et al*., 2016; Linington *et al*., 1988), along with an isotype control (rOCB-MS3-s1; Ctr-IgG; Brandle *et al*., 2016).

### Surgical procedure and in vivo imaging

Laminectomy surgeries were performed as previously described (Herwerth *et al*., 2016; Herwerth *et al*., 2022; Nikic *et al*., 2011). Mice were anesthetized via intraperitoneal injection of medetomidine (0.5 mg/kg), midazolam (5 mg/kg), and fentanyl (0.05 mg/kg), with additional doses administered as required. After performing a double dorsal laminectomy over lumbar segments L4-L5, mice were stabilized using compact spinal cord clamps (Davalos *et al*., 2008). The dura mater was punctured with a hypodermic needle and carefully removed to provide a dura-free imaging window. A 2% agarose well was constructed around the exposed area for application of artificial cerebrospinal fluid (aCSF; in mM: 148.2 NaCl, 3.0 KCl, 0.8 Na₂HPO₄, 0.2 NaH₂PO₄, 1.4 CaCl₂, 0.8 MgCl₂) or antibody/complement solutions. Recombinant antibodies (rAQP4-IgG vs rCtrl-IgG, 1.5 µg/mL; or MOG-IgG vs Ctr-IgG, 40 µg/mL) or heat-inactivated plasma (AQP4-IgG vs Ctrl-IgG, 150 µg/mL) were applied together with human complement source (healthy donor sera, 20% v/v of IgG/complement mix in aCSF) every 30 min for the first 2 hours, and subsequently every 60 min during the duration of the imaging session. In select experiments, ethidium homodimer-1 (EtHD, 1:500, 0.56 mg/mL stock; Invitrogen, Carlsbad, CA) was added to assess membrane integrity. For toxin-based astrocyte ablation experiments, intermedilysin (ILY, 200 ng/mL) was locally applied for 60 minutes, followed by aCSF washout, and imaging for up to 6 hours with aCSF. Our prior studies confirmed that neither phototoxicity nor transgenic labeling adversely affected glial cell health under these conditions (Herwerth *et al*., 2016; Herwerth *et al*., 2022).

In vivo imaging of the lumbar spinal cord was performed using a two-photon microscope (Olympus, Japan; FV1000-MPE or FVMPE-RS) equipped with femtosecond pulsed Ti:Sapphire Lasers (Mai Tai line, Newport/Spectra-Physics) using a water immersion objective (25x/1.05 N.A., Olympus). Excitation wavelengths were 770 nm (GFP/EtHD), 910 nm (GFP, GCaMP5g/tdTom, DsRed), or 950 nm (GFP/tdTom). Emission was filtered via a 690 nm short-pass dichroic mirror and separated using a G/R filter set (BA 495-540, BA 570-625) before detection with gallium arsenide phosphide (GaAsP) photomultiplier tubes. Time-lapse stacks were acquired every 10 minutes for glial morphology (up to 8 hours), or every 5 minutes for calcium imaging and microglia protrusion dynamics (up to 6 hours). Imaging parameters included up to 50 stacks with 1.5-2 µm z-steps, zooms of 1.0-2.0, and pixel sizes of 0.28-0.38 µm.

Astrocyte laser ablations were performed by individually targeting up to 15 cells within a region of interest. The center of each cell was marked with manually drawn ROIs, and ablations were carried out using a 950 nm laser at 80-120 mW for 0.5-1 minutes in “tornado” scan mode. No signs of additional photodamage (e.g., to non-targeted cells) were observed in either the ablated or control areas throughout the imaging period.

### Cell survival analysis

Cell death was defined as a ≥50% reduction in mean gray intensity or the appearance of morphological alterations: astrocyte disintegration or oligodendrocyte soma swelling with process blebbing/nuclear condensation. Given the clear distinction between control and NMO conditions, analyses were performed without blinding.

### GCaMP5g-based analysis of calcium signaling

Visually detectable calcium transients were identified from z-projections and confirmed in unprocessed 3D stacks. Mean gray values were measured in ImageJ/Fiji (Schindelin *et al*., 2012), with ROIs manually defined within each stack. GFP fluorescence intensity in each region of interest (ROI) was corrected by subtracting the mean gray value of a background ROI drawn in a cell-free region for each time frame and expressed as a percentage of the baseline (0 minutes) to quantify relative changes in intracellular calcium over time. Cells were considered to have elevated calcium if GCaMP5g fluorescence exceeded baseline by ≥50%. A steady-state intracellular calcium elevation was defined as GCaMP5g fluorescence ≥50% of baseline for at least 3 consecutive imaging time frames.

### Image processing

Image processing was performed using the open-source ImageJ/Fiji (Schindelin *et al*., 2012) and Adobe Creative Suite. Individual imaging channels were merged using pseudocolors for figure preparations, and brightness, contrast, and gamma were adjusted to enhance visibility in non-quantitative panels. Maximum intensity projections were used for display purposes, and time-lapse sequences were aligned using the StackReg plugin (Thevenaz *et al*., 1998).

### Data analysis

Sample sizes were chosen based on our previous in vivo imaging studies (Herwerth *et al*., 2016; Herwerth *et al*., 2022). Data were analyzed using Microsoft Excel (Microsoft Corporation, Redmond, WA) and the GraphPad Prism 10 software (GraphPad Software, San Diego, California, USA). The 50% times (t_50_) for cell survival and high calcium state were calculated by linear interpolation between the two nearest time points with survival values above and below 50%. Results are presented as mean ± SEM. Group comparisons were performed using non-parametric tests: Mann–Whitney U tests for two-group comparisons and Kruskal–Wallis tests for comparisons involving more than two groups, followed by appropriate post-hoc pairwise analyses when applicable. P-values < 0.05 were considered statistically significant and are denoted as follows: *P < 0.05; **P < 0.01; ***P < 0.001. Generative AI based on large language models (ChatGPT 5, Perplexity, Grammarly) was used to enhance writing and editing, but also to retrieve background information; all machine-generated text, information, and edits were checked and verified by the authors.

## Data availability

The datasets generated and analysed during the current study are available from the corresponding authors on reasonable request. Mouse lines can be requested under academic material transfer agreements from The Jackson Laboratories or from the laboratories that generated the lines, as indicated.

## Acknowledgements

We are grateful to D. Steinmetz and N. Budak for excellent animal husbandry and K. Kellermann for veterinary support. We thank Y. Hufnagel, M. Schetterer, and K. Wullimann for outstanding technical and administrative support. We also thank the labs of A. Waisman (University of Mainz) and B. Gao (NIH) for their generous provision of *MOG*-Cre deleter and huCD59 reporter mice, respectively. We acknowledge the SyNergy Microscale Hub for imaging support. Most funding for this work in T.M.’s and C.S.’s labs was provided by the Deutsche Forschungsgemeinschaft via TRR 274/1-2 2020, project B03 and Z01 – ID 408885537. Additional DFG-funded support for the project in T.M.’s lab came from grants Mi 694/9-1/2 A03 – ID 405358801, TRR 167/3 2025 – ID 259373024, and CRC1744, project A07 – ID 548585053. C.S.’s lab received additional funding for this project via STA1389/5-1 (individual research grant), STA1389/6-1, the DFG Priority Program 2395 “Local and peripheral drivers of microglia diversity and function” (project ID 500301720; STA1389/7-1/2), the DFG under Germany’s Excellence Strategy (EXC2067/1-390729940), the Hertie Foundation, and the National MS Society (USA). B.H., K.D., and T.M. receive support from the Munich Center for Systems Neurology (SyNergy EXC 2145 – ID 390857198). B.H. received funding by the European Union’s Horizon 2020 Research and Innovation Program (grant WISDOM, RIA 101137154). High-resolution microscopy was supported via a DFG instrumentation grant (INST95/1755-1 FUGG – ID 518284373). S.K., S.A., and K.E. are members of the Graduate School of Systemic Neurosciences (GSN).

## Author contributions

S.K., M.H., C.S., B.H., and T.M. conceptualized the project and experiments. S.K. and M.H. performed most of the investigation, including *in vivo* imaging and structural microscopy, and analyzed the data, but also managed mouse lines (supported by S.A. and K.E.). K.D., X.Q., and J.L.B. contributed reagents. M.H., B.H., C.S., and T.M. were responsible for funding acquisition; T.M. provided most of the animal and instrumentation resources. S.K and T.M. wrote the paper with input from all the authors.

## References

1. Akerboom J, Chen TW, Wardill TJ, Tian L, Marvin JS, Mutlu S, Calderon NC, Esposti F, Borghuis BG, Sun XR et al. (2012) Optimization of a GCaMP calcium indicator for neural activity imaging. J Neurosci 32: 13819–13840

2. Alberdi E, Sanchez-Gomez MV, Torre I, Domercq M, Perez-Samartin A, Perez-Cerda F, Matute C (2006) Activation of kainate receptors sensitizes oligodendrocytes to complement attack. J Neurosci 26: 3220–3228

3. Bacmeister CM, Barr HJ, McClain CR, Thornton MA, Nettles D, Welle CG, Hughes EG (2020) Motor learning promotes remyelination via new and surviving oligodendrocytes. Nat Neurosci 23: 819–831

4. Bayly-Jones C, Bubeck D, Dunstone MA (2017) The mystery behind membrane insertion: a review of the complement membrane attack complex. Philos Trans R Soc Lond B Biol Sci 372

5. Beltran E, Paunovic M, Gebert D, Cesur E, Jeitler M, Hoftberger R, Malotka J, Mader S, Kawakami N, Meinl E et al. (2021) Archeological neuroimmunology: resurrection of a pathogenic immune response from a historical case sheds light on human autoimmune encephalomyelitis and multiple sclerosis. Acta Neuropathol 141: 67–83

6. Bennett JL, Lam C, Kalluri SR, Saikali P, Bautista K, Dupree C, Glogowska M, Case D, Antel JP, Owens GP et al. (2009) Intrathecal pathogenic anti-aquaporin-4 antibodies in early neuromyelitis optica. Ann Neurol 66: 617–629

7. Berthele A, Levy M, Wingerchuk DM, Pittock SJ, Shang S, Kielhorn A, Royston M, Sabatella G, Palace J (2023) A single relapse induces worsening of disability and health-related quality of life in patients with neuromyelitis optica spectrum disorder. Front Neurol 14: 1099376

8. Black JA, Waxman SG (1988) The perinodal astrocyte. Glia 1: 169–183

9. Blaszczyk GJ, Mohammadnia A, Piscopo VEC, Sirois J, Cui QL, Yaqubi M, Durcan TM, Schneider R, Antel JP (2025) Pro-Inflammatory Molecules Implicated in Multiple Sclerosis Divert the Development of Human Oligodendrocyte Lineage Cells. Neurol Neuroimmunol Neuroinflamm 12: e200407

10. Bradl M, Lassmann H (2014) Experimental models of neuromyelitis optica. Brain Pathol 24: 74–82

11. Brandle SM, Obermeier B, Senel M, Bruder J, Mentele R, Khademi M, Olsson T, Tumani H, Kristoferitsch W, Lottspeich F et al. (2016) Distinct oligoclonal band antibodies in multiple sclerosis recognize ubiquitous self-proteins. Proc Natl Acad Sci U S A 113: 7864–7869

12. Breckwoldt MO, Pfister FM, Bradley PM, Marinkovic P, Williams PR, Brill MS, Plomer B, Schmalz A, St Clair DK, Naumann R et al. (2014) Multiparametric optical analysis of mitochondrial redox signals during neuronal physiology and pathology in vivo. Nat Med 20: 555–560

13. Brill MS, Lichtman JW, Thompson W, Zuo Y, Misgeld T (2011) Spatial constraints dictate glial territories at murine neuromuscular junctions. J Cell Biol 195: 293–305

14. Carnero Contentti E, Correale J (2021) Neuromyelitis optica spectrum disorders: from pathophysiology to therapeutic strategies. J Neuroinflammation 18: 208

15. Couves EC, Gardner S, Voisin TB, Bickel JK, Stansfeld PJ, Tate EW, Bubeck D (2023) Structural basis for membrane attack complex inhibition by CD59. Nat Commun 14: 890

16. Davalos D, Lee JK, Smith WB, Brinkman B, Ellisman MH, Zheng B, Akassoglou K (2008) Stable in vivo imaging of densely populated glia, axons and blood vessels in the mouse spinal cord using two-photon microscopy. J Neurosci Methods 169: 1–7

17. Doerflinger NH, Macklin WB, Popko B (2003) Inducible site-specific recombination in myelinating cells. Genesis 35: 63–72

18. Duan T, Smith AJ, Verkman AS (2018) Complement-dependent bystander injury to neurons in AQP4-IgG seropositive neuromyelitis optica. J Neuroinflammation 15: 294

19. Duan T, Verkman AS (2020) Experimental animal models of aquaporin-4-IgG-seropositive neuromyelitis optica spectrum disorders: progress and shortcomings. Brain Pathol 30: 13–25

20. Duncan ID, Radcliff AB, Heidari M, Kidd G, August BK, Wierenga LA (2018) The adult oligodendrocyte can participate in remyelination. Proc Natl Acad Sci U S A 115: E11807–E11816

21. Dunkelberger JR, Song WC (2010) Complement and its role in innate and adaptive immune responses. Cell Res 20: 34–50

22. Dutta DJ, Woo DH, Lee PR, Pajevic S, Bukalo O, Huffman WC, Wake H, Basser PJ, SheikhBahaei S, Lazarevic V et al. (2018) Regulation of myelin structure and conduction velocity by perinodal astrocytes. Proc Natl Acad Sci U S A 115: 11832–11837

23. Farkas I, Baranyi L, Ishikawa Y, Okada N, Bohata C, Budai D, Fukuda A, Imai M, Okada H (2002) CD59 blocks not only the insertion of C9 into MAC but inhibits ion channel formation by homologous C5b-8 as well as C5b-9. J Physiol 539: 537–545

24. Feng D, Dai S, Liu F, Ohtake Y, Zhou Z, Wang H, Zhang Y, Kearns A, Peng X, Zhu F et al. (2016) Cre-inducible human CD59 mediates rapid cell ablation after intermedilysin administration. J Clin Invest 126: 2321–2333

25. Fujihara K, Bennett JL, de Seze J, Haramura M, Kleiter I, Weinshenker BG, Kang D, Mughal T, Yamamura T (2020) Interleukin-6 in neuromyelitis optica spectrum disorder pathophysiology. Neurol Neuroimmunol Neuroinflamm 7

26. Gee JM, Smith NA, Fernandez FR, Economo MN, Brunert D, Rothermel M, Morris SC, Talbot A, Palumbos S, Ichida JM et al. (2014) Imaging activity in neurons and glia with a Polr2a-based and cre-dependent GCaMP5G-IRES-tdTomato reporter mouse. Neuron 83: 1058–1072

27. Gregorian C, Nakashima J, Le Belle J, Ohab J, Kim R, Liu A, Smith KB, Groszer M, Garcia AD, Sofroniew MV et al. (2009) Pten deletion in adult neural stem/progenitor cells enhances constitutive neurogenesis. J Neurosci 29: 1874–1886

28. Guo Y, Weigand SD, Popescu BF, Lennon VA, Parisi JE, Pittock SJ, Parks NE, Clardy SL, Howe CL, Lucchinetti CF (2017) Pathogenic implications of cerebrospinal fluid barrier pathology in neuromyelitis optica. Acta Neuropathol 133: 597–612

29. Herwerth M, Kalluri SR, Srivastava R, Kleele T, Kenet S, Illes Z, Merkler D, Bennett JL, Misgeld T, Hemmer B (2016) In vivo imaging reveals rapid astrocyte depletion and axon damage in a model of neuromyelitis optica-related pathology. Ann Neurol 79: 794–805

30. Herwerth M, Kenet S, Schifferer M, Winkler A, Weber M, Snaidero N, Wang M, Lohrberg M, Bennett JL, Stadelmann C et al. (2022) A new form of axonal pathology in a spinal model of neuromyelitis optica. Brain 145: 1726–1742

31. Hinson SR, Pittock SJ, Lucchinetti CF, Roemer SF, Fryer JP, Kryzer TJ, Lennon VA (2007) Pathogenic potential of IgG binding to water channel extracellular domain in neuromyelitis optica. Neurology 69: 2221–2231

32. Hoftberger R, Guo Y, Flanagan EP, Lopez-Chiriboga AS, Endmayr V, Hochmeister S, Joldic D, Pittock SJ, Tillema JM, Gorman M et al. (2020) The pathology of central nervous system inflammatory demyelinating disease accompanying myelin oligodendrocyte glycoprotein autoantibody. Acta Neuropathol 139: 875–892

33. Hovelmeyer N, Hao Z, Kranidioti K, Kassiotis G, Buch T, Frommer F, von Hoch L, Kramer D, Minichiello L, Kollias G et al. (2005) Apoptosis of oligodendrocytes via Fas and TNF-R1 is a key event in the induction of experimental autoimmune encephalomyelitis. J Immunol 175: 5875–5884

34. Hu W, Ferris SP, Tweten RK, Wu G, Radaeva S, Gao B, Bronson RT, Halperin JA, Qin X (2008) Rapid conditional targeted ablation of cells expressing human CD59 in transgenic mice by intermedilysin. Nat Med 14: 98–103

35. Jakel S, Agirre E, Mendanha Falcao A, van Bruggen D, Lee KW, Knuesel I, Malhotra D, Ffrench-Constant C, Williams A, Castelo-Branco G (2019) Altered human oligodendrocyte heterogeneity in multiple sclerosis. Nature 566: 543–547

36. Jung S, Aliberti J, Graemmel P, Sunshine MJ, Kreutzberg GW, Sher A, Littman DR (2000) Analysis of fractalkine receptor CX(3)CR1 function by targeted deletion and green fluorescent protein reporter gene insertion. Mol Cell Biol 20: 4106–4114

37. Kimberley FC, Sivasankar B, Paul Morgan B (2007) Alternative roles for CD59. Mol Immunol 44: 73–81

38. Kumpfel T, Giglhuber K, Aktas O, Ayzenberg I, Bellmann-Strobl J, Haussler V, Havla J, Hellwig K, Hummert MW, Jarius S et al. (2024) Update on the diagnosis and treatment of neuromyelitis optica spectrum disorders (NMOSD) - revised recommendations of the Neuromyelitis Optica Study Group (NEMOS). Part II: Attack therapy and long-term management. J Neurol 271: 141–176

39. Lennon VA, Kryzer TJ, Pittock SJ, Verkman AS, Hinson SR (2005) IgG marker of optic-spinal multiple sclerosis binds to the aquaporin-4 water channel. J Exp Med 202: 473–477

40. Lennon VA, Wingerchuk DM, Kryzer TJ, Pittock SJ, Lucchinetti CF, Fujihara K, Nakashima I, Weinshenker BG (2004) A serum autoantibody marker of neuromyelitis optica: distinction from multiple sclerosis. Lancet 364: 2106–2112

41. Linington C, Bradl M, Lassmann H, Brunner C, Vass K (1988) Augmentation of demyelination in rat acute allergic encephalomyelitis by circulating mouse monoclonal antibodies directed against a myelin/oligodendrocyte glycoprotein. Am J Pathol 130: 443–454

42. Liu Y, Given KS, Harlow DE, Matschulat AM, Macklin WB, Bennett JL, Owens GP (2017) Myelin-specific multiple sclerosis antibodies cause complement-dependent oligodendrocyte loss and demyelination. Acta Neuropathol Commun 5: 25

43. Liu Y, Harlow DE, Given KS, Owens GP, Macklin WB, Bennett JL (2016) Variable sensitivity to complement-dependent cytotoxicity in murine models of neuromyelitis optica. J Neuroinflammation 13: 301

44. Lucchinetti CF, Guo Y, Popescu BF, Fujihara K, Itoyama Y, Misu T (2014) The pathology of an autoimmune astrocytopathy: lessons learned from neuromyelitis optica. Brain Pathol 24: 83–97

45. Lucchinetti CF, Mandler RN, McGavern D, Bruck W, Gleich G, Ransohoff RM, Trebst C, Weinshenker B, Wingerchuk D, Parisi JE et al. (2002) A role for humoral mechanisms in the pathogenesis of Devic’s neuromyelitis optica. Brain 125: 1450–1461

46. Madisen L, Zwingman TA, Sunkin SM, Oh SW, Zariwala HA, Gu H, Ng LL, Palmiter RD, Hawrylycz MJ, Jones AR et al. (2010) A robust and high-throughput Cre reporting and characterization system for the whole mouse brain. Nat Neurosci 13: 133–140

47. Mallon BS, Shick HE, Kidd GJ, Macklin WB (2002) Proteolipid promoter activity distinguishes two populations of NG2-positive cells throughout neonatal cortical development. J Neurosci 22: 876–885

48. Marignier R, Hacohen Y, Cobo-Calvo A, Probstel AK, Aktas O, Alexopoulos H, Amato MP, Asgari N, Banwell B, Bennett J et al. (2021) Myelin-oligodendrocyte glycoprotein antibody-associated disease. Lancet Neurol 20: 762–772

49. Marignier R, Nicolle A, Watrin C, Touret M, Cavagna S, Varrin-Doyer M, Cavillon G, Rogemond V, Confavreux C, Honnorat J et al. (2010) Oligodendrocytes are damaged by neuromyelitis optica immunoglobulin G via astrocyte injury. Brain 133: 2578–2591

50. Marignier R, Villacieros-Alvarez J, Espejo C, Arrambide G, Fissolo N, Gutierrez L, Dinoto A, Mulero P, Rubio-Flores L, Nieto P et al. (2025) Assessment of neuronal and glial serum biomarkers in myelin oligodendrocyte glycoprotein antibody-associated disease: the MULTIMOGAD study. J Neurol Neurosurg Psychiatry 96: 884–892

51. Masaki K (2015) Early disruption of glial communication via connexin gap junction in multiple sclerosis, Balo’s disease and neuromyelitis optica. Neuropathology 35: 469–480

52. Matute C (2007) Interaction between glutamate signalling and immune attack in damaging oligodendrocytes. Neuron Glia Biol 3: 281–285

53. Mezydlo A, Treiber N, Ullrich Gavilanes EM, Eichenseer K, Ancau M, Wens A, Ares Carral C, Schifferer M, Snaidero N, Misgeld T et al. (2023) Remyelination by surviving oligodendrocytes is inefficient in the inflamed mammalian cortex. Neuron 111: 1748–1759 e1748

54. Misu T, Fujihara K, Kakita A, Konno H, Nakamura M, Watanabe S, Takahashi T, Nakashima I, Takahashi H, Itoyama Y (2007) Loss of aquaporin 4 in lesions of neuromyelitis optica: distinction from multiple sclerosis. Brain 130: 1224–1234

55. Misu T, Hoftberger R, Fujihara K, Wimmer I, Takai Y, Nishiyama S, Nakashima I, Konno H, Bradl M, Garzuly F et al. (2013) Presence of six different lesion types suggests diverse mechanisms of tissue injury in neuromyelitis optica. Acta Neuropathol 125: 815–827

56. Nagamune H, Ohnishi C, Katsuura A, Fushitani K, Whiley RA, Tsuji A, Matsuda Y (1996) Intermedilysin, a novel cytotoxin specific for human cells secreted by Streptococcus intermedius UNS46 isolated from a human liver abscess. Infect Immun 64: 3093–3100

57. Neely SA, Williamson JM, Klingseisen A, Zoupi L, Early JJ, Williams A, Lyons DA (2022) New oligodendrocytes exhibit more abundant and accurate myelin regeneration than those that survive demyelination. Nat Neurosci 25: 415–420

58. Nicaise C, Marneffe C, Bouchat J, Gilloteaux J (2019) Osmotic Demyelination: From an Oligodendrocyte to an Astrocyte Perspective. Int J Mol Sci 20

59. Nikic I, Merkler D, Sorbara C, Brinkoetter M, Kreutzfeldt M, Bareyre FM, Bruck W, Bishop D, Misgeld T, Kerschensteiner M (2011) A reversible form of axon damage in experimental autoimmune encephalomyelitis and multiple sclerosis. Nat Med 17: 495–499

60. Nimmerjahn A, Kirchhoff F, Helmchen F (2005) Resting microglial cells are highly dynamic surveillants of brain parenchyma in vivo. Science 308: 1314–1318

61. Owens GP, Fellin TJ, Matschulat A, Salas V, Schaller KL, Given KS, Ritchie AM, Navarro A, Blauth K, Hughes EG et al. (2023) Pathogenic myelin-specific antibodies in multiple sclerosis target conformational proteolipid protein 1-anchored membrane domains. J Clin Invest 133

62. Park CC, Shin ML, Simard JM (1997) The complement membrane attack complex and the bystander effect in cerebral vasospasm. J Neurosurg 87: 294–300

63. Parratt JD, Prineas JW (2010) Neuromyelitis optica: a demyelinating disease characterized by acute destruction and regeneration of perivascular astrocytes. Mult Scler 16: 1156–1172

64. Peschl P, Schanda K, Zeka B, Given K, Bohm D, Ruprecht K, Saiz A, Lutterotti A, Rostasy K, Hoftberger R et al. (2017) Human antibodies against the myelin oligodendrocyte glycoprotein can cause complement-dependent demyelination. J Neuroinflammation 14: 208

65. Piddlesden SJ, Morgan BP (1993) Killing of rat glial cells by complement: deficiency of the rat analogue of CD59 is the cause of oligodendrocyte susceptibility to lysis. J Neuroimmunol 48: 169–175

66. Ratelade J, Asavapanumas N, Ritchie AM, Wemlinger S, Bennett JL, Verkman AS (2013) Involvement of antibody-dependent cell-mediated cytotoxicity in inflammatory demyelination in a mouse model of neuromyelitis optica. Acta Neuropathol 126: 699–709

67. Richard C, Ruiz A, Cavagna S, Bigotte M, Vukusic S, Masaki K, Suenaga T, Kira JI, Giraudon P, Marignier R (2020) Connexins in neuromyelitis optica: a link between astrocytopathy and demyelination. Brain 143: 2721–2732

68. Roemer SF, Parisi JE, Lennon VA, Benarroch EE, Lassmann H, Bruck W, Mandler RN, Weinshenker BG, Pittock SJ, Wingerchuk DM et al. (2007) Pattern-specific loss of aquaporin-4 immunoreactivity distinguishes neuromyelitis optica from multiple sclerosis. Brain 130: 1194–1205

69. Rollins SA, Zhao J, Ninomiya H, Sims PJ (1991) Inhibition of homologous complement by CD59 is mediated by a species-selective recognition conferred through binding to C8 within C5b-8 or C9 within C5b-9. J Immunol 146: 2345–2351

70. Romanelli E, Merkler D, Mezydlo A, Weil MT, Weber MS, Nikic I, Potz S, Meinl E, Matznick FE, Kreutzfeldt M et al. (2016) Myelinosome formation represents an early stage of oligodendrocyte damage in multiple sclerosis and its animal model. Nat Commun 7: 13275

71. Saadoun S, Waters P, Bell BA, Vincent A, Verkman AS, Papadopoulos MC (2010) Intra-cerebral injection of neuromyelitis optica immunoglobulin G and human complement produces neuromyelitis optica lesions in mice. Brain 133: 349–361

72. Saadoun S, Waters P, Owens GP, Bennett JL, Vincent A, Papadopoulos MC (2014) Neuromyelitis optica MOG-IgG causes reversible lesions in mouse brain. Acta Neuropathol Commun 2: 35

73. Schindelin J, Arganda-Carreras I, Frise E, Kaynig V, Longair M, Pietzsch T, Preibisch S, Rueden C, Saalfeld S, Schmid B et al. (2012) Fiji: an open-source platform for biological-image analysis. Nat Methods 9: 676–682

74. Scolding NJ, Morgan BP, Compston DA (1998) The expression of complement regulatory proteins by adult human oligodendrocytes. J Neuroimmunol 84: 69–75

75. Sharma R, Fischer MT, Bauer J, Felts PA, Smith KJ, Misu T, Fujihara K, Bradl M, Lassmann H (2010) Inflammation induced by innate immunity in the central nervous system leads to primary astrocyte dysfunction followed by demyelination. Acta Neuropathol 120: 223–236

76. Snaidero N, Schifferer M, Mezydlo A, Zalc B, Kerschensteiner M, Misgeld T (2020) Myelin replacement triggered by single-cell demyelination in mouse cortex. Nat Commun 11: 4901

77. Takai Y, Hametner S, Riedl C, Misu T, Takahashi T, Suzuki H, Chihara N, Watanabe M, Miyahara H, Yoshida M et al. (2026) Characteristic patterns of complement deposition in NMOSD, MOGAD, and MS. Acta Neuropathol 151: 13

78. Takai Y, Misu T, Kaneko K, Chihara N, Narikawa K, Tsuchida S, Nishida H, Komori T, Seki M, Komatsu T et al. (2020) Myelin oligodendrocyte glycoprotein antibody-associated disease: an immunopathological study. Brain 143: 1431–1446

79. Takizawa H, Takahashi K, Murakami T, Okada N, Okada H (1992) Species-specific restriction of complement by HRF20 (CD59) generated by cDNA transfection. Eur J Immunol 22: 1943–1946

80. Thevenaz P, Ruttimann UE, Unser M (1998) A pyramid approach to subpixel registration based on intensity. IEEE Trans Image Process 7: 27–41

81. Tradtrantip L, Yao X, Su T, Smith AJ, Verkman AS (2017) Bystander mechanism for complement-initiated early oligodendrocyte injury in neuromyelitis optica. Acta Neuropathol 134: 35–44

82. Triantafilou K, Hughes TR, Triantafilou M, Morgan BP (2013) The complement membrane attack complex triggers intracellular Ca2+ fluxes leading to NLRP3 inflammasome activation. J Cell Sci 126: 2903–2913

83. Uzawa A, Oertel FC, Mori M, Paul F, Kuwabara S (2024) NMOSD and MOGAD: an evolving disease spectrum. Nat Rev Neurol 20: 602–619

84. Vidal-Itriago A, Radford RAW, Aramideh JA, Maurel C, Scherer NM, Don EK, Lee A, Chung RS, Graeber MB, Morsch M (2022) Microglia morphophysiological diversity and its implications for the CNS. Front Immunol 13: 997786

85. Wang Z, Guo W, Liu Y, Gong Y, Ding X, Shi K, Thome R, Zhang GX, Shi FD, Yan Y (2017) Low expression of complement inhibitory protein CD59 contributes to humoral autoimmunity against astrocytes. Brain Behav Immun 65: 173–182

86. Weil MT, Mobius W, Winkler A, Ruhwedel T, Wrzos C, Romanelli E, Bennett JL, Enz L, Goebels N, Nave KA et al. (2016) Loss of Myelin Basic Protein Function Triggers Myelin Breakdown in Models of Demyelinating Diseases. Cell Rep 16: 314–322

87. Williams PR, Marincu BN, Sorbara CD, Mahler CF, Schumacher AM, Griesbeck O, Kerschensteiner M, Misgeld T (2014) A recoverable state of axon injury persists for hours after spinal cord contusion in vivo. Nat Commun 5: 5683

88. Wing MG, Zajicek J, Seilly DJ, Compston DA, Lachmann PJ (1992) Oligodendrocytes lack glycolipid anchored proteins which protect them against complement lysis. Restoration of resistance to lysis by incorporation of CD59. Immunology 76: 140–145

89. Wingerchuk DM, Banwell B, Bennett JL, Cabre P, Carroll W, Chitnis T, de Seze J, Fujihara K, Greenberg B, Jacob A et al. (2015) International consensus diagnostic criteria for neuromyelitis optica spectrum disorders. Neurology 85: 177–189

90. Wingerchuk DM, Lucchinetti CF (2022) Neuromyelitis Optica Spectrum Disorder. N Engl J Med 387: 631–639

91. Winkler A, Wrzos C, Haberl M, Weil MT, Gao M, Mobius W, Odoardi F, Thal DR, Chang M, Opdenakker G et al. (2021) Blood-brain barrier resealing in neuromyelitis optica occurs independently of astrocyte regeneration. J Clin Invest 131

92. Witte ME, Schumacher AM, Mahler CF, Bewersdorf JP, Lehmitz J, Scheiter A, Sanchez P, Williams PR, Griesbeck O, Naumann R et al. (2019) Calcium Influx through Plasma-Membrane Nanoruptures Drives Axon Degeneration in a Model of Multiple Sclerosis. Neuron 101: 615–624 e615

93. Wrzos C, Winkler A, Metz I, Kayser DM, Thal DR, Wegner C, Bruck W, Nessler S, Bennett JL, Stadelmann C (2014) Early loss of oligodendrocytes in human and experimental neuromyelitis optica lesions. Acta Neuropathol 127: 523–538

94. Wu G, Chen T, Shahsafaei A, Hu W, Bronson RT, Shi GP, Halperin JA, Aktas H, Qin X (2010) Complement regulator CD59 protects against angiotensin II-induced abdominal aortic aneurysms in mice. Circulation 121: 1338–1346

95. Wu G, Hu W, Shahsafaei A, Song W, Dobarro M, Sukhova GK, Bronson RR, Shi GP, Rother RP, Halperin JA et al. (2009) Complement regulator CD59 protects against atherosclerosis by restricting the formation of complement membrane attack complex. Circ Res 104: 550–558

96. Xie CB, Jane-Wit D, Pober JS (2020) Complement Membrane Attack Complex: New Roles, Mechanisms of Action, and Therapeutic Targets. Am J Pathol 190: 1138–1150

97. Yang Y, Vidensky S, Jin L, Jie C, Lorenzini I, Frankl M, Rothstein JD (2011) Molecular comparison of GLT1+ and ALDH1L1+ astrocytes in vivo in astroglial reporter mice. Glia 59: 200–207

98. Yao X, Verkman AS (2017a) Complement regulator CD59 prevents peripheral organ injury in rats made seropositive for neuromyelitis optica immunoglobulin G. Acta Neuropathol Commun 5: 57

99. Yao X, Verkman AS (2017b) Marked central nervous system pathology in CD59 knockout rats following passive transfer of Neuromyelitis optica immunoglobulin G. Acta Neuropathol Commun 5: 15

100. Yu J, Dong S, Rushmere NK, Morgan BP, Abagyan R, Tomlinson S (1997) Mapping the regions of the complement inhibitor CD59 responsible for its species selective activity. Biochemistry 36: 9423–9428

101. Zhu X, Bergles DE, Nishiyama A (2008) NG2 cells generate both oligodendrocytes and gray matter astrocytes. Development 135: 145–157

