## Supplementary material for "Early demyelination by off-target complement injury in a mouse model of neuromyelitis optica"

### Supplementary Information

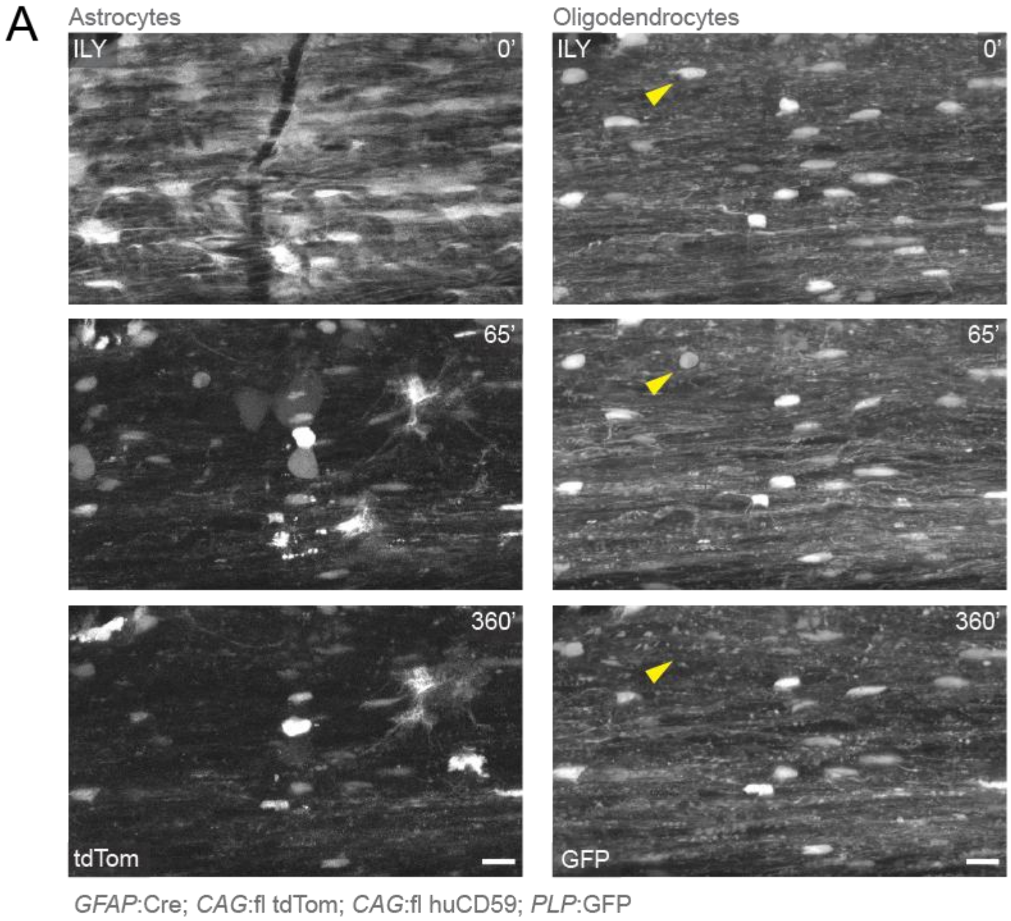


**Supplementary Figure 1. Selective, toxin-based astrocyte depletion does not induce oligodendrocyte injury**

**A.** Local spinal application of intermedilysin toxin (200ng/mL) for 1-hour resulted in marked depletion of astrocytes. The same region was imaged for up to 6 hours following ILY exposure. Despite profound astrocytic loss, oligodendrocyte somata remain largely unaffected with the exception of a single cell exhibiting swelling with spheroid soma and subsequent cell death (yellow arrowheads). Scale bar 20µm.

**Supplementary Video 1**

In vivo spinal cord two-photon imaging in *ALDH1L1*:GFP; *PLP*:CreERT x *CAG*:fl tdTom mouse over an 8-hour time-lapse, corresponding to images shown in **Figure 1A, B**. Local application of AQP4-IgG and complement induced experimental lesions observed with global, lytic astrocyte death (green) followed by oligodendrocyte injury (magenta). Scale bar 40 μm.

**Supplementary Video 2**

Magnified (3.5x) in vivo two-photon time-lapse of oligodendrocytes showing early signs of injury along processes with beadings (arrowheads) and subsequently progressed to a swollen phenotype with the appearance of a spheroid soma. *ALDH1L1*:GFP; *PLP*:CreERT x *CAG*:fl tdTom mouse, corresponding to AQP4-IgG treatment images in **Figure 1A**. Scale bar 20 μm.

**Supplementary Video 3**

In vivo spinal cord two-photon imaging in *GFAP*:Cre; CAG-fl-PCGCaMP5g-IRES-tdTom mouse over an 3-hour time-lapse, corresponding to images shown in **Figure 3A**. Local application of r-AQP4-IgG and complement induced calcium signals (orange) in astrocytes (green). Scale bar 30 μm.

**Supplementary Video 4**

In vivo spinal cord two-photon imaging in *PLP*:CreERT; CAG-fl-PCGCaMP5g-IRES-tdTom mouse over an 4-hour time-lapse, corresponding to images shown in **Figure 3C**. Local application of r-AQP4-IgG and complement induced calcium signals (orange) in oligodendrocytes. Scale bar 30 μm.
